# Pooled CRISPRi screening of the cyanobacterium *Synechocystis* sp. PCC 6803 for enhanced growth, tolerance, and chemical production

**DOI:** 10.1101/823534

**Authors:** Lun Yao, Kiyan Shabestary, Sara M. Björk, Johannes Asplund-Samuelsson, Haakan N. Joensson, Michael Jahn, Elton P. Hudson

## Abstract

We developed an inducible CRISPRi gene repression library in the cyanobacterium *Synechocystis* sp. PCC 6803, where all annotated genes are targeted for repression. We used the library to estimate gene fitness in multiple conditions. The library revealed several mutants with increased specific growth rates (up to 17%), and transcriptomics of these mutants revealed common upregulation of genes within photosynthetic electron flow. We challenged the library with L-lactate stress to find more tolerant mutants. Repression of the peroxiredoxin Bcp2 increased growth rate by 49% in the presence of 0.1 M L-lactate. Finally, the library was transformed into a L-lactate-secreting strain, and droplet microfluidics sorting of top producers enriched sgRNAs targeting nutrient assimilation, redox modulation, and cyclic-electron flow. Several clones showed increased productivity in batch cultivations (up to 75%). In some cases, tolerance or productivity was enhanced by partial repression of essential genes, which are difficult to access by transposon insertion.

## Introduction

Cyanobacteria are model organisms for photosynthetic electron flow, photorespiration, and the circadian clock^1–3^. In addition to their massive ecological importance, biotechnological applications of cyanobacteria have been proposed, such as microbial cell factories, where metabolism is engineered to synthesize chemicals from CO_2_ using energy derived from light^4–6^. This widespread interest in cyanobacteria brings a need for system-wide analysis of gene essentiality and function. For many bacteria, the transposon mutagenesis library has been used to identify *loss-of-function* or *gain-of-function* mutants. New variants of transposon mutagenesis tag the transposon insertion site with a barcode, allowing mutants to be tracked using nextgeneration sequencing (NGS)^7, 8^. However, transposon insertions can have sequence bias, and mutants with knockouts of essential genes are lost at the time of library creation.

An alternative to transposon mutant libraries are pooled CRISPRi libraries for targeted gene repression, where unique single-guide RNA (sgRNA) genes are pooled and transformed into the strain of interest. Since the protospacer region of sgRNAs is small enough (~ 20 nt) to be sequenced by NGS, it can serve as a barcode and allow monitoring of the abundance of each clone in the library. CRISPRi libraries have been applied to screen gene essentiality and diverse phenotypes including morphology, solvent tolerance, and phage resistance^9–11^. Inducible CRISPRi has been developed for several cyanobacteria strains^12, 13^, but pooled sgRNA libraries have not been exploited. A transposon library in the cyanobacterium *Synechococcus elongatus* PCC 7942 (hereafter *Synechococcus* PCC 7942) was used to map essential genes in both photoautotrophic and diurnal growth^14, 15^.

Here we report the construction and use of an inducible CRISPRi library of > 10,000 clones for the model cyanobacterium *Synechocystis sp*. PCC 6803 (hereafter *Synechocystis*). We tracked the composition of the library during growth in multiple conditions, including different light regimes and in the presence of the organic acid L-lactate. We also screened the library for enhanced L-lactate production using droplet microfluidics (Fig. 1). In addition to providing gene fitness scores for almost all open reading frames and non-coding RNAs in *Synechocystis*, our results give new insights into how to engineer cyanobacteria metabolism for industrial use. We show that the growth rate of *Synechocystis* can be improved through several knockdowns. Further, we verify previous computational predictions that altering ATP/NADPH balance can improve bioproduction in cyanobacteria. By screening both growth and productivity, this platform could yield mutants that trade biomass formation for increased biochemical production. The data for all competition experiments performed with the sgRNA library can be accessed through an interactive web application at https://m-jahn.shinyapps.io/ShinyLib/.

**Figure 1.**
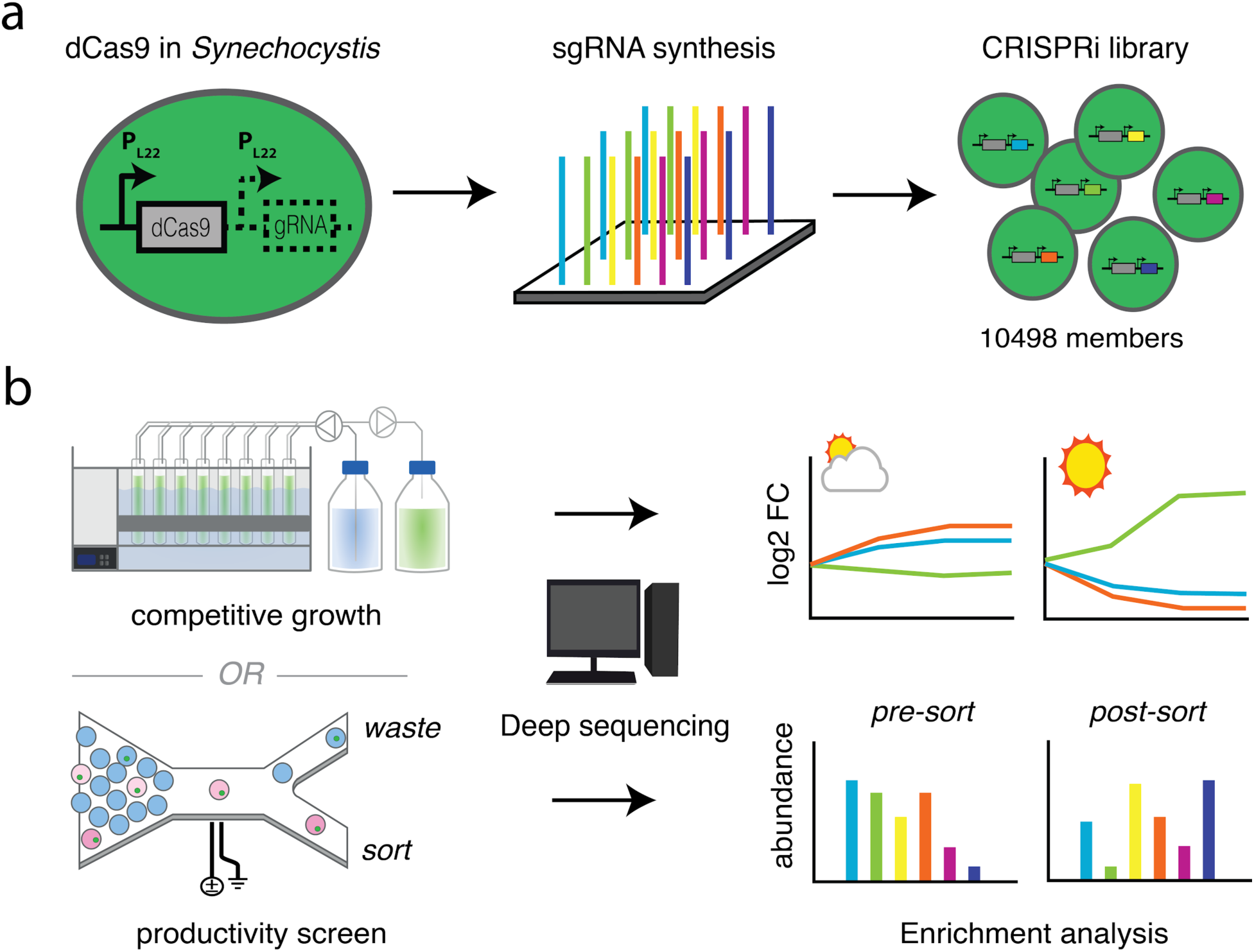
Workflow for CRISPRi screening in *Synechocystis*. **a** Creating the CRISPRi library. An inducible promoter P_L22_ is used in *Synechocystis*. **b** The fitness or productivity of each library member is assessed by counting mutants in a bioreactor cultivation or in a sorted population using next-generation sequencing.

## Results

### Gene fitness during photoautotrophic growth

We designed a CRISPRi library for repression of each annotated gene (ORF, ncRNA, and asRNA) in *Synechocystis*^16^. Two sgRNAs were designed for each target (Methods, Supplemental Data 1, Supplemental Data 2). The resulting 10498 sgRNA sequences were synthesized, pooled, and cloned into a genomic integration vector, and the pooled sgRNAs were transformed into a *Synechocystis* strain containing a genome-integrated, anhydrotetracycline (aTc)-inducible dCas9 cassette^13^. The resultant *Synechocystis* CRISPRi library was characterized by NGS of the sgRNA sequences; 10496 sgRNAs were present.

We cultivated the library in light-limited turbidostats under two constant-light conditions, 100 μmol photons m^-2^ s^-1^ (L100) and 300 μmol photons m^-2^ s^-1^ (L300) and a light-dark diurnal condition (LD, 0 to 300 μmol photons m^-2^ s^-1^), each with supplemented 1% CO_2_ and in 4 replicates. Samples for NGS were taken periodically over 32 days, on average corresponding to 16 and 32 generations for the L100 and LD cultivations (μ ~ 0.03 h^-1^), and L300 cultivations (μ ~ 0.07 h^-1^), respectively (Supplemental Data 3). The abundance of 7119 sgRNAs (clones) targeting ORFs were quantified in each condition and averaged across the 4 replicates (Supplemental Data 4). The sgRNAs were grouped into 5 clusters based on their depletion patterns (Fig. 2a). In total, 1998 sgRNAs (28.1%) had a significant depletion in at least one condition, indicating these target genes have some contribution to cell fitness (Fig. 2b). Cluster 1 (239 sgRNAs) contained sgRNAs that were depleted in induced and non-induced cultivations, revealing weak background leakage of dCas9 and particular sensitivity to changes in abundance of these genes. Cluster 2 (370 sgRNAs) was enriched in sgRNAs that were quickly depleted in all growth conditions. Cluster 3 (880 sgRNAs) contained sgRNAs that were more rapidly depleted in L300 than L100. Clusters 4 (503 sgRNAs) and 5 (5121 sgRNAs) contained sgRNAs that were depleted slowly or not at all. A gene-ontology (GO term) enrichment analysis showed that clusters 1 and 2 were highly enriched for GO terms related to essential cellular processes such as photosynthesis, carbon fixation, and translation (Fig. 2c). Cyanobacteria invest most of their resources into synthesizing proteins from these groups^17, 18^. Interestingly, nearly all genes with a low fitness score are also among proteins found by Jahn *et al*. to be regulated with growth rate (Supplemental Fig. 1). Cluster 3 was enriched for a more diverse set of GO terms (*e.g*. secondary metabolites, cell membrane, and transport) and cluster 4 was enriched in only a few GO terms (nucleic acid metabolism and TCA cycle).

**Figure 2.**
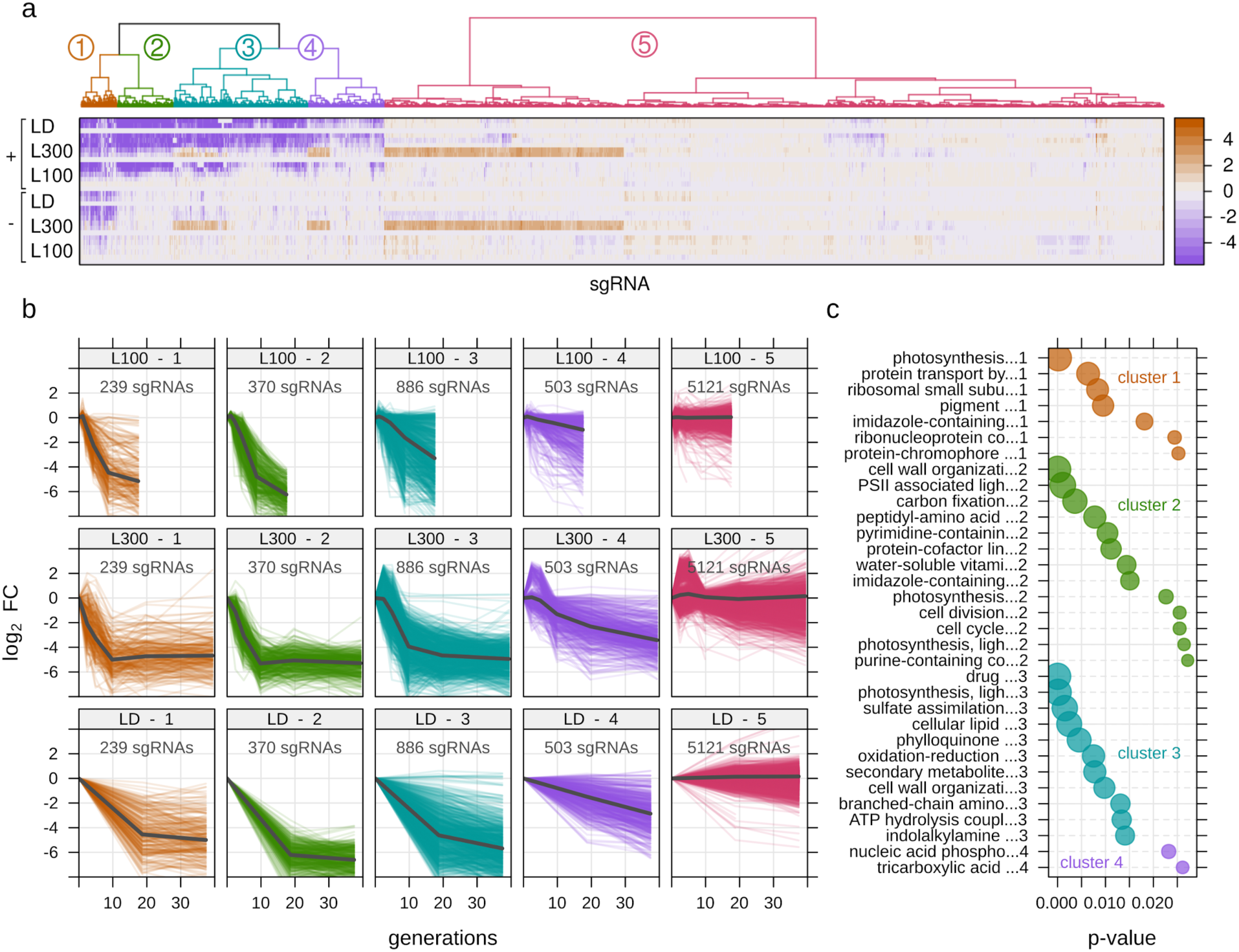
Enrichment or depletion of CRISPRi library mutants during photoautotrophic conditions. **a** Clustered sgRNAs by similarity of log_2_ fold-change over time. Rows represent time-points sampled after induction. (L100 - light with 100 μmol m^-2^ s^-1^, L300 - light with 300 μmol m^-2^ s^-1^, LD - light-dark cycle). All log_2_ fold-change values were calculated from averages of 4 replicate cultivations. Symbols: ‘+’ induced, ‘-’ non-induced. **b** Log_2_ fold change of individual sgRNAs over the course of each cultivation. Experiment run time was normalized to number of cell generations estimated from population growth rate. **c** Enriched gene ontology (GO) terms for the four clusters (1-4) showing sgRNA depletion; p-value - Fisher’s exact test with elimination (see Methods).

The depletion data revealed instances of both rigidity and plasticity in *Synechocystis* central carbon metabolism. We examined genes in central carbon metabolism for effects of CRISPRi repression (Supplemental Fig. 2). Repression of malic enzyme (*slr0721*) and pyruvate kinase (*pyk2*, *sll1275*) both resulted in a growth defect (Supplemental Data 4), even though pyruvate kinase is predicted to carry little flux^19^. Both routes to pyruvate are therefore important. Within the Calvin cycle, two energetically equivalent routes for synthesis of fructose-6-phosphate (F6P) have been considered: through class-I fructose-bisphosphate (FBP) aldolase (*fda,slr0943*) or flux through transaldolase (*talB, slr1793*)^20^. The repression of either of these genes had no effect on photoautotrophic growth, though the class-II FBP/SBP aldolase gene (*cbbA, sll0018*), which is needed to synthesize sedoheptulose bisphosphate (SBP) for both routes, was essential. While these results do not definitively show which route is favored under photoautotrophic conditions, they suggest that flux can operate through either route if the other is perturbed. This is in contrast to *Synechococcus* PCC 7942, where the sole FBP aldolase gene was found to be essential for photoautotrophic growth, and transaldolase was not^14^.

We next compared fitness scores for sgRNAs in each growth condition (see Methods for calculation). Fitness scores were generally lower in the L300 condition than in L100, even when normalized to number of cell generations (Fig. 3a). One explanation could be a higher protein turnover rate in the higher growth condition, as reported for *Lactococcus lactis*^21^. A second explanation is that faster growing cells are more prone to perturbation of enzyme levels, due to higher enzyme saturation. We found 42 genes that were beneficial for growth in L300 but neutral in L100 (Fig. 3b). This set was almost exclusively in cluster 3 and included genes related to DNA repair, proteome and redox homeostasis, and subunits of PSI and the NADPH dehydrogenase NDH-1 (Fig. 3c, Supplemental Data 5). Energy dissipation and photo-stress response are thus particularly important for growth at high light. There were 25 genes with differential fitness between L100 and LD, and only 4 between L300 and LD (Supplemental Fig. 3 and 4). This is far fewer than in *Synechococcus* PCC 7942 where more than 100 genes were found with a transposon library to be important for LD growth^15^. The discrepancy could be attributed to the CRISPRi library comprising partial repressions and not gene knockouts, and that many of the genes with a fitness effect in LD are involved in alleviating light stresses experienced at dawn, making them also important for the L300 condition.

**Figure 3.**
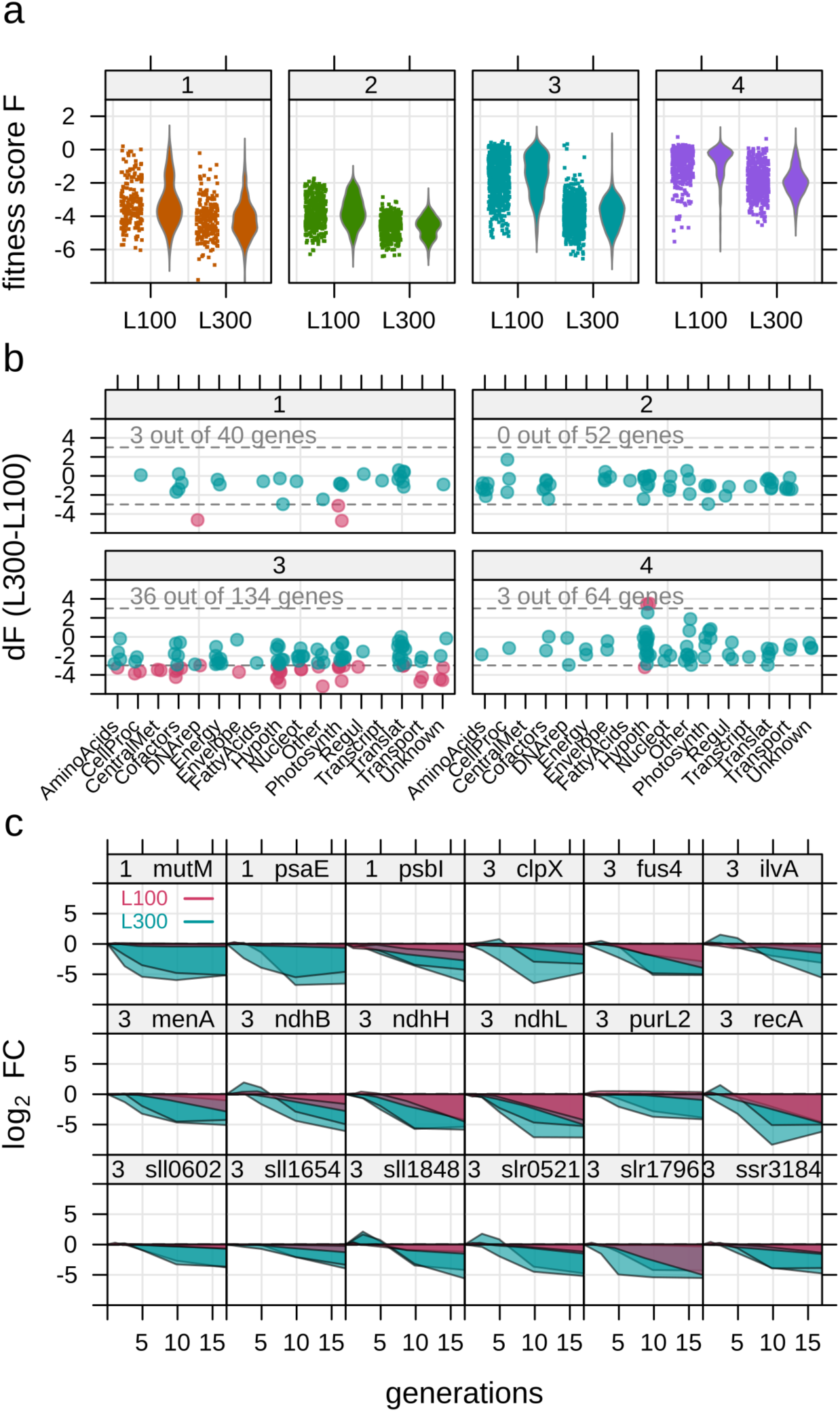
Genes with condition-dependent fitness in different light conditions. **a** Distribution of fitness score *F* for all sgRNAs in clusters 1 to 4. Fitness score indicates the degree of enrichment (positive) or depletion (negative) of an sgRNA, normalized to cell generations. Comparison between two light conditions (L100 - light with 100 μmol m^-2^ s^-1^ and L300 - light with 300 μmol m^-2^ s^-1^) shows sgRNAs are on average more rapidly depleted under L300 independent of number of cell generations. **b** Difference in *F* (*dF*) between L100 and L300, for genes with both sgRNAs in the same cluster. **c** Time courses of both sgRNAs for the top 18 genes with highest *dF* in L100 and L300.

### Mutants with a growth advantage

Quantification of the library population over time in a turbidostat allows calculation of the maximum specific growth rate μ of each mutant (Methods). Growth rate estimates allow for more intuitive interpretation of mutant fitness and highlights that library clones are repression mutants, not total knockouts. For example, repression of Calvin cycle genes often caused significant reduction of μ (*e.g. prk* −40%, *cbbA* −95%), while repression of most photorespiration-related genes did not (*e.g. glcD1* −10%, *glcD2* −10%) (Supplemental Fig. 2). Many clones showed higher growth rates than the population average, notably *pmgA* (mean increase for L100 and L300 +17%) and *slr1916* (mean increase +13%) (Fig. 4a, Supplemental Data 6). PmgA is a regulator involved in the high-light response in *Synechocystis* and a *pmgA* knockout mutant has an inability to reduce PSI content in high light, resulting in more efficient photosynthesis, but also glucose sensitivity^22^. A *slr1916* mutant was previously identified from a small *Synechocystis* transposon library on the basis of altered fluorescence kinetics and also shows a higher PSI content at high light and glucose sensitivity^23^. Notably, other mutants isolated in the study of Ozaki *et al*. were not enriched in our library, indicating that altered PSI/PSII alone does not ensure faster growth. *slr1916* was annotated based on homology as *menH*, an esterase in the phylloquinone pathway^24^. However, repression of the 8 other phylloquinone pathway genes resulted in strong growth defects, suggesting *slr1916* is not a key enzyme in this pathway (Supplemental Fig. 5). Furthermore, *slr1916* is localized to the plasma membrane^25^, though its proposed substrates are soluble. Three additional sgRNA clones showed slight but significant increases in growth rate (+5%): *sll1969*, an annotated triacylglycerol lipase in the same operon as *pmgA; slr1340*, an uncharacterized gene with encoding a predicted acetyl-transferase, and *ssl2982*, encoding the ω subunit of RNA polymerase^26^. While the function of *slr1340* is not known, acetyl-transferases can have many roles in bacteria, including post-translational regulation of proteins through lysine acetylation^27^.

**Figure 4.**
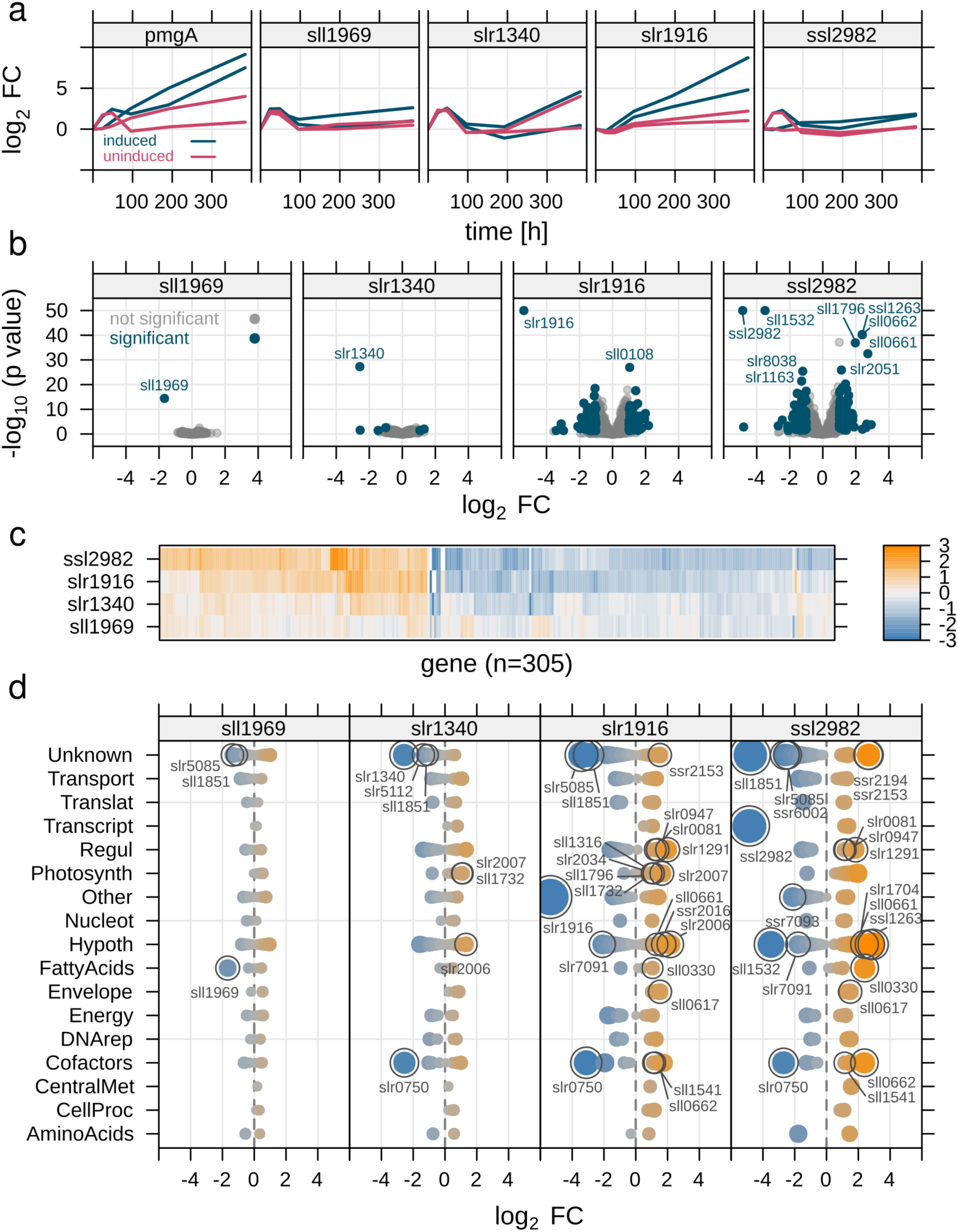
Transcriptomics of faster growing mutants. **a** Enrichment of faster growing sgRNA mutants in library competition experiment at L300 (300 μmol photons m^-2^ s^-1^), log_2_ fold change (log_2_ FC) of read count over time (n = 2). **b** Volcano plot of log_2_ FC for reconstructed mutants, compared to the control strain (sgRNA-NT0), against adjusted p-value for each gene. Grey - non-significantly different genes, blue - significantly different genes (threshold: negative log_10_ p-value ≥ 2; absolute log_2_ FC ≥ 1). The p-value for three sgRNA mutants was outside the plotting region and was restricted to −log_10_ of 50 for visibility. **c** Heat map of all 305 genes (log_2_ FC) that were significantly different in at least one of the four mutants, clustered into two different groups based on similarity of gene expression. **d** Significantly different genes sorted by Cyanobase pathways. Genes of particular interest were highlighted (see text for details). Blue and orange indicates negative and positive log_2_ FC, respectively. Size of symbols increases with increasing absolute log_2_ FC.

In an attempt to uncover common regulations among these mutants, we reconstructed *slr1916, sll1969, slr1340, and ssl2982* sgRNA mutants and collected transcriptomics after CRISPRi induction. A *pmgA* clone was not included as microarray data from a *pmgA* knockout was reported previously^28^. From the RNA-Seq data, *slr1916* and *ssl2982* mutants had 143 and 248 differential transcripts compared to a control strain (sgRNA-NT0, with no target site in *Synechocystis* genome) (Fig. 4b, Supplemental Data 7). A weak transcriptomic response of *slr1340* and *sll1969* mutants could be due to a lower repression efficiency in these clones (log_2_ FC of target genes was −2.57 and −1.67, respectively, compared to −5.37 and −4.85 for *slr1916* and *ssl2982*). Clustering genes based on similarity of expression changes revealed that the same set of genes was affected for all mutants, but with stronger effects in *slr1916* and *ssl2982* (Fig. 4c). The upregulated genes are involved in electron transport (*fed7*), and include components of the NDH-1 complex (*ndhD1, ndhD2*, and *ssr2016*) and carotenoid biosynthesis (*crtQ* and *crtZ*) (Fig. 4d). An upregulation of carotenoid biosynthesis suggests increased PSII repair. Cyanobacterial NDH-1 complexes participate in cyclic electron flow around PSI, respiratory electron flow, and CO_2_ uptake^29–31^. An increase in PSI activity at high light can dissipate pressure in the electron transport chain; a similar phenomenon was reported to contribute to high-light tolerance and faster growth in *Synechococcus elongatus* UTEX 2973 compared to its close relative *Synechococcus* PCC 7942^32^. Interestingly, the high-light responsive transcription factor RpaB (*slr0947*, regulator of phycobilisome association B) was also up-regulated in multiple mutants. The extensive regulon of RpaB includes likely repressor activity of linear electron transport at high light (*e.g*. subunits of PSII and Cyt-b_6_f), and activation activity of photoprotection and cyclic electron flow (*e.g. ftsH*, *ssr2016*)^33^. Genes that were down-regulated in all four mutants were *sll1851*, a small non-annotated gene, and *chlN* (*slr0750*), a subunit of the light-independent operative protochlorophyllide oxidoreductase (LI-POR).

### Gene fitness in the presence of L-lactate

The CRISPRi library is also beneficial for finding stress tolerance phenotypes, which are generally difficult to rationally engineer^34^. The commodity chemical L-lactate has been produced in several cyanobacteria but still at relatively low titers (15 mM^35^. The tolerance of *Synechocystis* to L-lactate is approximately 0.1 M (9 g/L), at which specific growth rate is reduced by 50% (L100 condition, Supplemental Data 3). To identify mutants with increased tolerance to L-lactate, we cultivated the library in turbidostats with 0.1 M sodium L-lactate (pH-adjusted) and sampled periodically for NGS (0, 16 and 32 d). We found 78 sgRNAs that were enriched during the L-lactate cultivation, but not in a NaCl control cultivation (Fig. 5a). Eight genes were enriched with both sgRNAs (Fig. 5b). Curiously, 19 of the enriched sgRNAs (24%) targeted genes in amino acid metabolism and protein biosynthesis, including multiple aminoacid tRNA synthetases (*argS, aspS, asnS, glyS, gltX, metS*) (Supplemental Fig. 6). Many of these clones had growth defect in the absence of L-lactate, which supports the general phenomenon that slow-growing microbes are more stress tolerant^36^. However, a more amino acid-specific effect is possible, as reduced biosynthesis could result in an accumulation of some amino acids, which have been implicated in stress resistance in bacteria^37^.

**Figure 5.**
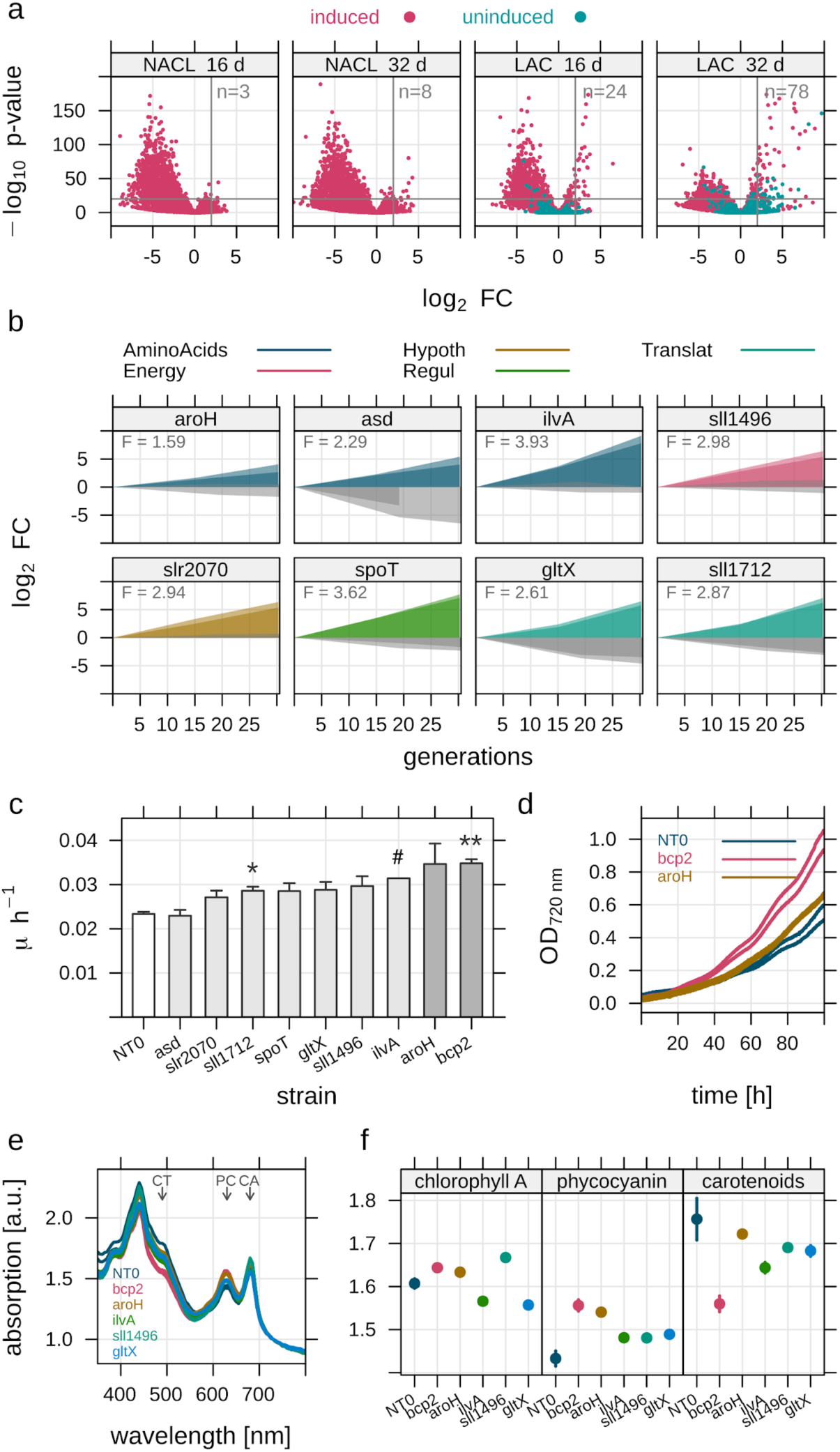
Identification of sgRNA clones with improved L-lactate tolerance. **a** Volcano plot showing enrichment of 78 sgRNAs during cultivation with added 0.1 M sodium L-lactate after 32 d (LAC), (threshold: log_2_ FC ≥ 2, −log_10_ p-value ≥ 20). Only 8 sgRNAs were enriched in a 0.1 M NaCl control cultivation (NACL). **b** Eight genes where both sgRNAs were enriched above average in 0.1 M L-lactate cultivation (coloured) but not NaCl cultivation (grey). Color indicates association to cyanobase pathway of the repressed gene. *F* - mean fitness score of two sgRNAs. **c** Mean growth rate μ (h^-1^) of selected re-constructed knock-down strains (n = 2) over the first 80 h of batch cultivation with L-lactate added to 0.1 M. Symbols show significance between the control strain (NT0) and mutants using student’s t-test. * - p-value ≤ 0.05, ** - p-value ≤ 0.01, # - only 1 replicate was used for *ilvA* mutant. **d** Example of growth advantage in batch culture (0.1 M added L-lactate) of sgRNA mutants *bcp2* and *aroH* over the control strain (NT0). **e** Absorption spectra of five fastest growing sgRNA mutants and NT0 control strain in 0.1 M L-lactate. Arrows mark absorption maxima for pigments. CT - carotenoids, PC - phycocyanin, CA - chlorophyll A. **f** Relative pigment absorption for carotenoids, phycocyanin and chlorophyll A for the five fastest growing mutants and NT0 control strain in presence of 0.1 M L-lactate.

We re-constructed nine of the repression clones and validated growth improvement in batch cultures with added L-lactate (Fig. 5c, Supplemental Fig. 7). We tested L-lactate consumption of these mutants and found that none consumed L-lactate over a 48-hour period. The *bcp2* mutant (*bacterioferritin co-migratory protein*, a peroxiredoxin) had the most significant tolerance improvement (49% increase in μ, p = 0.006, student’s t-test, Fig. 5d). The mechanism for increased tolerance in the *bcp2* mutant is not known. However, thioredoxins can mediate direct reduction of cysteines on transcription factors, including the master RpaB regulator in *Synechocystis*^38^. The absorbance spectra of all L-lactate-tolerant mutants showed increased chlorophyll A and phycocyanin absorption in the presence of L-lactate, suggesting higher photosynthetic activity (Fig. 5e-f). Relative concentration of carotenoids, pigments related to the light stress response, was reduced in L-lactate tolerant mutants.

We could also identify clones with growth negatively affected by L-lactate but not by NaCl (Supplemental Fig. 8). These targeted genes could thus be candidates for overexpression to improve tolerance. Most prominent were an antibiotic resistance gene (*zam, sll1910*), two nucleases, the protease *clpX* (*sll0535*), cytochrome M (*cytM, sll1245*) that may act to dissipate excess electrons^39^, and *sepF* (*slr2073*), an inhibitor of cell division.

### Screening the CRISPRi library for L-lactate productivity

The CRISPRi library can be linked to screens other than growth, and we next sought to find mutant clones that had increased productivity of L-lactate. To screen the library for a secreted product, we used droplet encapsulation and microfluidic sorting^40–42^. The sgRNA pool and inducible *dCas9* cassette were cloned into a *Synechocystis* strain containing lactate dehydrogenase from *Lactococcus lactis*^43^. The library was grown in shake-flasks, gene repression was induced, and samples were taken after 36 and 66 h to assay for L-lactate productivity (Methods, Supplemental Fig. 9). For each sample, approximately 180,000 cell-containing droplets were screened and 36,000 droplets were sorted. Cells could not be reliably recovered on agar plates after sorting, so we performed PCR of the sgRNA region directly from sorted droplets, followed by NGS library preparation and quantification. Approximately 2500 unique clones were detected with confidence (> 32 reads) from the sorted samples (Supplemental Data 8). Due to the small number of sorted cells and high variability between replicates, it was not possible to determine the significance of enrichment for each clone in the sorted populations. To assess which clones produced more L-lactate than average, we used instead the criterium that a clone had to be among the top 10% of enriched clones in at least two of the four sorted samples (Methods). This criterium returned 528 clones, 359 harbored sgRNAs targeting ORFs and 169 harbored sgRNAs targeted to ncRNAs (Supplemental Data 8).

Though nearly half of the enriched clones targeted genes of unknown function, it is possible to derive engineering strategies from the annotated targets (Table 1). Restriction of nutrient uptake or assimilation is known to trigger large alterations in carbon flux in cyanobacteria^44–46^. The enriched targets include glutamate dehydrogenase (*gdhA*), glutamate synthase (*gltB*), and glutamine synthase (*glnA*), repression of these would restrict NH3 assimilation^47^, as well as nitrogen (*nrtD2*) and phosphorous (*pstC*) transporters. CRISPRi repression of *glnA* was recently demonstrated in *Synechococcus* sp. PCC 7002 to improve lactate productivity 2-fold^12^. A second engineering strategy is suggested by target genes in electron transport and energy metabolism. Computational and experimental studies have predicted that efficient production of some biochemicals would require lowering the ATP/NADPH ratio in the cyanobacteria cell^48–50^. The enriched targets *ssr2016* and *ndhD2* are both involved in ATP-generating cyclic electron flow around PSI^30, 51^. *SdhB* is a subunit of succinate dehydrogenase directly contributing to respiration^52^. Finally, direct alteration of carbon flux was also apparent among enriched clones. Repression of citrate synthase (*gltA*) was shown to increase specific productivity of n-butanol in *Synechocystis*^53^. Repression of phosphoketolase (*slr0453*) may increase L-lactate productivity by eliminating a possible bypass of pyruvate^54, 55^.

**Table 1:** Selection of *Synechocystis* mutants enriched by droplet sorting for high L-lactate productivity. Mutants were identified from a CRISPRi library by fluorescence-based droplet sorting. Appearance is the frequency that each clone was ranked in the top 10% by enrichment factor in the high-lactate sorting gate (out of 4 samples). Impaired effect on growth is denoted when the sgRNA clone is depleted in L300 dataset after 4 generations (p_adj_ < 0.005). Asterisk denotes sgRNA mutants selected for reconstruction and validation in this study. Full table in Supplemental Data 8.

| sgRNA | Appearances | Effect on growth | Function |
| --- | --- | --- | --- |
| gdhA_25* | 4 | none | glutamate dehydrogenase |
| ssr2016_62* | 3 | impaired | ferredoxin:plastoquinone reductase, cyclic electron flow |
| gltA_41* | 3 | impaired | citrate synthase |
| gltB_3* | 3 | none | glutamate synthase |
| ndhD2_15 | 3 | none | NDH-1 complex subunit, cyclic electron flow |
| sdhB_26* | 2 | none | succinate dehydrogenase |
| Entry85_13* | 2 | none | atpA (ATP synthase) asRNA |
| pstC_70 | 2 | none | phosphate transporter |
| pntB_19 | 2 | none | transhydrogenase subunit |

We re-constructed six clones for validation of L-lactate productivity, representing targets within nutrient uptake, carbon flux, and redox and energy generation (Table 1). In a first screen, we cultivated the clones in shake-flasks. Cultures were induced for gene repression 2 days prior to inoculation. The *gltA* strain had a significantly higher L-lactate titer than the control strain (Fig. 6a). The effect was enhanced when titers were normalized to cell density, indicating a redirection of carbon flux from biomass formation to product^53^. Two of the potentially redox-altered mutants (*sdhB, ssr2016*) had slight increases (20%) in L-lactate titer, though not statistically significant. We next tested these mutants in a ‘photonfluxostat’ reactor, where light intensity was gradually increased with cell density to insure a fixed light dosage per cell^56^. This cultivation mode was expected to activate alternative electron flow reactions for an extended time, so as to amplify any effects of repressing these on L-lactate productivity. In photonfluxostat mode, the *sdhB* and *ssr2016* clones had higher L-lactate titers than the control strain (Fig. 6b). We note that titers and specific productivities were lower for all strains in the photonfluxostat mode than in shake-flasks, which could be due to altered gas transfer or different perceived light intensities.

**Figure 6.**
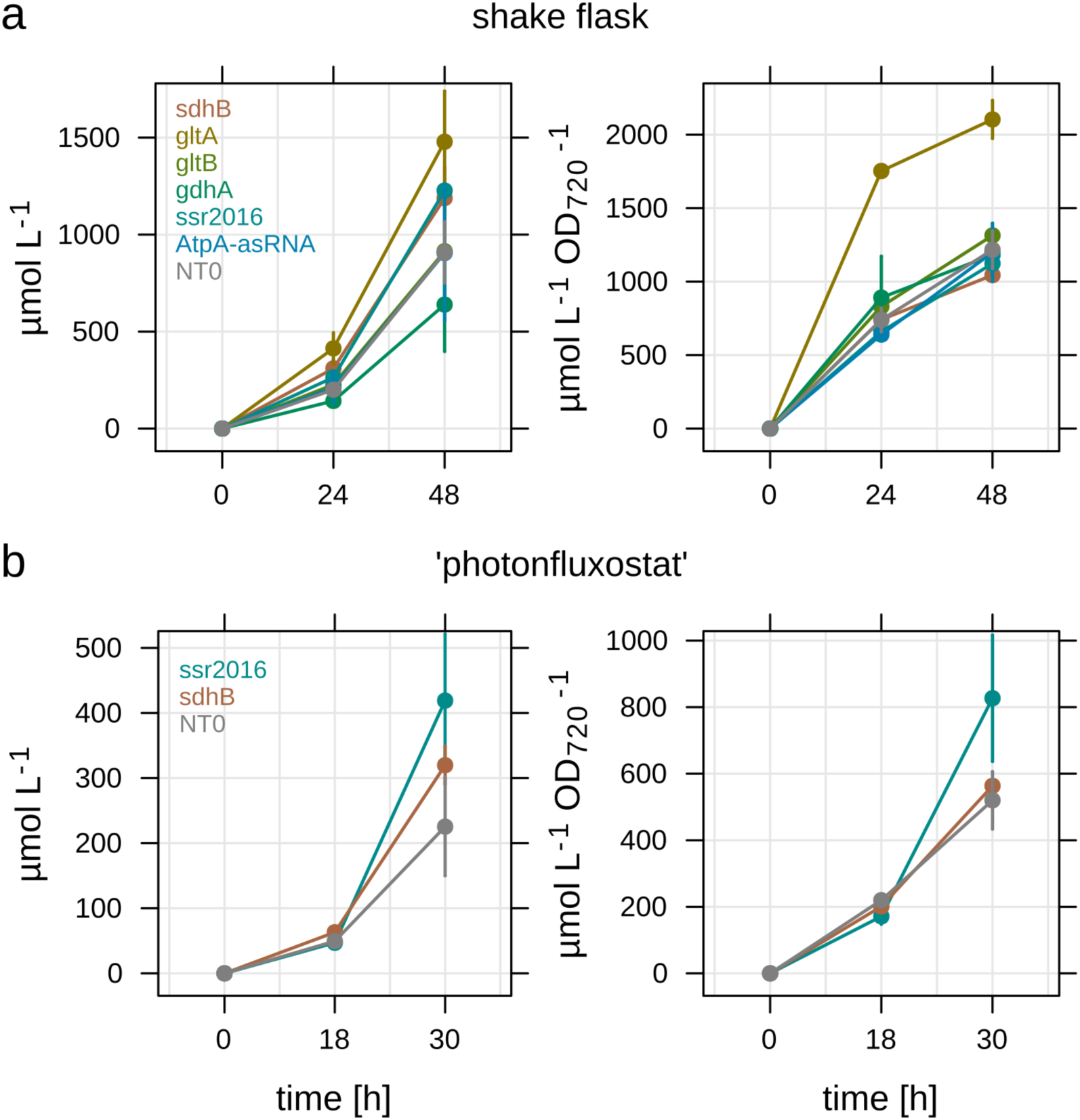
SgRNA mutants with increased L-lactate production. Mutants with potentially increased L-lactate productivity were discovered using a fluorescence-activated droplet sorting assay. Six selected enriched mutants were re-constructed, cultivated, and product titer determined in μmol per liter, and μmol per liter per biomass (OD_720 nm_). **a** Productivity of enriched mutants cultivated in axenic shake-flask cultures (n = 2). L-lactate concentration was measured after 24 and 48 hours. NT0 - control sgRNA with no target site in *Synechocystis* genome. **b** A subset of two mutants (excluding the known over-producer *gltA*) was cultivated in axenic ‘photonfluxostat’ cultures (light intensity proportional to cell density, 1000 μmol photons m^-2^ s^-1^ OD_720 nm_^-1^, n = 2). L-lactate concentration was measured after 18 and 30 hours.

## Discussion

We have primarily used the CRISPRi repression library to identify and validate mutants with improved industrial phenotypes. By providing fitness contributions of all genes in a certain condition, the CRISPRi library reveals patterns in how cyanobacteria solve certain physiological challenges. There were multiple genetic avenues for enhancing the growth rate of *Synechocystis*, though they converged on a similar transcriptome that upregulates photosystem repair as well as electron-transport around PSI. This is in line with a recent modeling study, which showed that the rate of ATP and NADPH generation exerted the most control over carbon fixation rates in *Synechocystis*^57^. Interestingly, four out of five sgRNA targets in top faster-growing strains are potentially regulator proteins (*pmgA*, *slr1916*, *ssl2982*, *slr1340*). A recent report showed that fast growth could be accessed in the cyanobacterium *Synechococcus* PCC 7942 with point mutations in energy-generating enzymes such as ATP synthase, but the full fast-growth phenotype also required mutations in the master transcription factor RpaA that drastically altered the transcriptome^58^. The downregulation of *chlN*, a subunit of the light-independent operative protochlorophyllide oxidoreductase (LI-POR), in three of the four faster-growing clones is intriguing as chlorophyll is connected to photosystem biosynthesis^59^. The LI-POR enzyme appears to be fully dispensable during photoautotrophic growth, but mutation of the *chlL* subunit reduced both growth and PSI content during light-activated heterotrophic cultivations^59^. Transcripts of the other subunits of LI-POR, *chlB* and *chlL*, were not affected in the faster-growing mutants.

Care should be taken when using results from CRISPRi libraries to determine gene essentiality. We used 2 sgRNAs targeting each ORF; recent studies have shown that more are needed to ensure statistical significance when assessing gene essentiality^11^. For approximately 50% of the genes with high fitness scores in our data, one sgRNA was significantly depleted from the library and the other was not (Supplemental Fig. 10). There are several lines of evidence which suggest that this discrepancy is due to weak binding of one of the sgRNAs (a false negative) instead of off-target binding of an sgRNA to an essential gene elsewhere (false positive). First, the repression efficiency of sgRNAs can range from 50-99% (Fig. 4b and Yao *et al*.^13^) so partial repression of an essential gene may not elicit a phenotype. Second, genes where at least one sgRNA clone is depleted show 87% overlap (287/329) with genes predicted to be essential by Flux Balance Analysis of a *Synechocystis* genome-scale model^60^ (Supplemental Data 9). There is also a 75% agreement in our calculated fitness scores for genes in central carbon metabolism to their orthologs in *Synechoccocus* PCC 7942, the latter determined by Tn-Seq^14^ (Supplemental Data 10). Third, off-target binding by dCas9 was shown to be problematic for strongly expressed dCas9^61^, while our genome-integrated *dCas9* was driven by a weak promoter^62^. However, dCas9 is potent, as we observed gene repression for some targets even in the absence of the inducer (Cluster 1 in Fig. 2). Finely-graded dCas9 expression may require native, metal-sensitive promoters^63, 64^ or addition of translational-level control such as riboswitches^65^.

The CRISPRi library has a unique advantage over gene knockout libraries for engineering bioproduction in that the level of essential genes can be titrated, allowing perturbation of the core metabolic network^66^. Cultures where a ‘metabolic switch’ shifts metabolism away from growth can be more productive^53, 67, 68^. By coupling CRISPRi library to a fluorescence assay, we were able to screen thousands of potential knockdowns for increased productivity, giving a first test of many metabolic engineering strategies previously proposed by computational modeling. However, several limitations were apparent in the library-droplet microfluidics workflow. The dynamic nature of CRISPRi repression could cloud beneficial effects if the timing of sorting after induction is not optimized. The 4 h incubation period where cells secrete L-lactate generates high single-cell variability and is dependent on cell shading, though we attempted to counter this with a high screening depth. A short incubation period was chosen to prevent saturation of the fluorescence assay^69^. Larger droplet sizes would allow for longer incubation times. In batch validation, not all of the enriched clones from the droplet sorting gave higher L-lactate titer and there were apparent tradeoffs between productivity and carbon partitioning in some clones. Therefore, we propose that the enriched clones from a CRISPRi library are most useful for creating guiding principles for engineering, or that repression of individual targets could be combined to get more significant increases in productivity.

It is easy to envision expanding an inducible CRISPRi library to other screens. Since the two fastest-growing mutants found here were also glucose sensitive, it could be worthwhile to add a screen for photomixotrophic growth, which results in flux through alternative glycolytic pathways^70^. Small transposon libraries (300 mutants) have previously been used to identify genes required for phototaxis by screening for colony smearing^71^; a CRISPRi library could greatly accelerate identification of genes involved in mobility. A fluorescence-activated cell sorting (FACS) screen for cell fluorescence could be useful in mapping the photobleaching program during nutrient limitation and screens for recovery ability from starvation could identify carbon metabolism pathways involved in resuscitation^72^. Such a screen would be a particularly fertile application for an inducible CRISPRi library since central carbon metabolism mutants are not lost during initial library creation. Finally, libraries limited to a subset of target genes would be amenable to multiplexing of gene repression^73^.

## Methods

### Genetic constructs and cloning of sgRNA library

The *Synechocystis* base strain containing the *tetR*_P_L22__*dCas9* expression cassette in the *psbA1* locus (spectinomycin resistance) was described previously^13^. The sgRNA library oligos were synthesized on a 12K chip by CustomArray Inc., USA (see Supplemental Data 1 for a list of all sgRNA sequences). The pooled oligos were cloned into a genome targeting vector (locus *slr0397*^74^, kanamycin resistance) containing the P_L22_ promoter by Golden Gate assembly. NEB 10-beta Competent *E. coli* cells (New England BioLabs) were used for library transformation and a total number of 1,200,000 *E. coli* colonies were obtained. All colonies were collected, resuspended in LB, pooled together and then cultivated overnight in LB for plasmid extraction. Natural transformation was used to create sgRNA libraries in *Synechocystis*^75^, where approximately 300,000 colonies were obtained. Colonies were collected, resuspended in fresh BG-11 and pooled. The pooled *Synechocystis* library was stored at −80°C. When re-creating specific sgRNAs, we used overlap-extension PCR of a template sgRNA as described previously^13^. All primers are listed in Supplemental Table 1. A L-lactate-secreting *Synechocystis* strain was created by cloning the *ldh* gene with L39R substitution from *Lactococcus lactis* under the P*trc* promoter^43^. This gene construct was inserted into the genome at locus *slr0168* with a chloramphenicol resistance cassette. This strain was subsequently transformed with the *tetR*_P_L22__*dCas9* cassette and sgRNA library as described above, resulting in a L-lactate-producing *Synechocystis* sgRNA library.

### sgRNA library design

Two sgRNAs were designed for each open reading frame (ORF) and non-coding RNA in the *Synechocystis* genome (Reference genome NCBI NC_00091). ORF sequence annotations were obtained from NCBI (downloaded on 18.03.2016). Locations of non-coding RNAs were from^76^. An in-house Python script was used to create protospacer sequences as close to the transcription start site (TSS) or translation start codon (ATG) as possible, within the following criteria: Target regions were required to be within 500 bp from the known TSS or within 75% of total gene length, absence of G_6_ and T_4_, and GC content between 25% and 75%. Target sequences were searched according to the pattern 5’-CCN[20-25 bases])T-3’. The 5’-CCN ensured a 5’-NGG-3’ PAM site on the coding strand, and 3’ T was to ensure binding of the 5’ end of the sgRNA, known to have an A when transcribed from promoter P_L22_ in *Synechocystis*^62^.

For each target fitting these criteria, potential off-target regions in the genome were then identified. An off-target binding site defined as having fewer than two mismatches in the PAM-proximal 17 bp region of the proposed sgRNA. Both NGG and NAG PAMs and both strands were considered. Then all sgRNA candidates for a gene, the two sgRNAs with the least off-targets were selected. If possible, sgRNAs were selected that were at least 10 bp apart from each other.

### Cultivation in photobioreactor turbidostats

The *Synechocystis* sgRNA library was cultivated in an 8-tube photobioreactor (Multi-Cultivator MC-1000-OD, Photon System Instruments, Drasov, CZ). The system was customized to perform turbidostat cultivation as described previously^17^. Reactors (65 mL) were bubbled with 1% v/v CO_2_ in air at 2.5 mL/min, and light intensity was controlled by a computer program. The OD_720 nm_ and OD_680 nm_ were measured every 15 min. The turbidity set point was OD_720 nm_ = 0.2 and 22 mL fresh BG-11 was added to dilute the culture once the set point was exceeded. Anhydrotetrocycline inducer was added to 500 ng/uL to the culture to induce *dCas9* expression, and to the reserve BG-11 media used for dilution. For LD cultures, the light regime (12 h light - 12 h dark) followed a sinusoidal function with maximum light intensity at 300 μmol m^-2^ s^-1^ (L (t) = 300 ⋅ sin (π/43200⋅ t) where t = cultivation time in seconds). For turbidostat cultivations with added L-lactate, light intensity was 100 μmol m^-2^ s^-1^ and either 100 mM sodium L-lactate or 100 mM NaCl was added to the BG-11 before inoculation and medium pH was adjusted to 7.8. All turbidostat cultivations described in this work were performed as 4 independent replicates. To sample for NGS, 15 mL of culture volume was harvested by centrifugation (5,000 g for 5 min, 25°C). Cells were collected and stored at −20°C. Batch cultivations for growth rate or L-lactate quantification of selected sgRNA mutants were performed in the same conditions as turbidostat cultures, but with light at 300 μmol photons m^-2^ s^-1^. Mutants were pre-cultivated in BG-11 with antibiotics (25 μg/mL spectinomycin, 25 μg/mL kanamycin) and aTc inducer (added to 500 ng/mL) in a climatic chamber (Percival Climatics SE-1100 with 100 μE/s/m^2^ illumination and 1% v/v CO_2_) for 3 days, and then inoculated into the photobioreactor to OD_730 nm_ = 0.05 before growth measurements began.

### Next-generation sequencing of sgRNAs

A two-step PCR procedure was carried out for NGS library preparation using sgRNA library plasmid from *E.coli* or genomic DNA from *Synechocystis* library as template. Genomic DNA was extracted from *Synechocystis* cell pellets (15 mL culture at OD_720 nm_ 0.2) using GeneJET Genomic DNA Purification Kit (Thermo Fisher Scientific). 1^st^ step PCR was performed using the primer pair LUYA593/LUYA594, which amplify sgRNAs from the plasmid or genome and add adaptors for NGS. The PCR product was analyzed on Agilent 2100 Bioanalyzer (Agilent Technologies) and gel purified using GeneJET Gel Extraction Kit (Thermo Fisher Scientific). The purified DNA was used as template to perform the 2^nd^ step PCR using NEBNext Multiplex Oligos for Illumina (Dual Index Primers Set 1) (New England Biolabs), followed by Bioanalyzer analysis and gel purification. Purified DNA was quantified using Qubit fluorometer 2.0 (Thermo Fisher Scientific) and then pooled. The NGS was performed on Illumina NextSeq 500 system using NextSeq 500/550 High Output v2 kit (75 cycles). In a typical NGS run, 40 samples were analyzed simultaneously, providing 50 to 100 reads per sgRNA per sample (Supplemental Fig. 11). We used sickle 1.33 (https://github.com/najoshi/sickle) to trim (75 nt) and clean reads. A custom python script was used to assign and count reads to each sgRNA.

### Fitness score calculation

The gradual, sgRNA-mediated depletion of clones from the library allows estimation of the contribution to cellular fitness for each individual gene. Here, we defined the fitness *F* of a mutant as the area under the curve (AUC) for log_2_ fold change sgRNA abundance (log_2_ FC) at a number of generations (n_gen_) since induction, normalized by maximum generations.

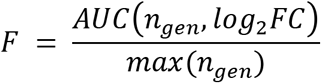

Differential fitness between for example two light conditions L300, L100 was calculated as = *ΔF* = *F*_*L*300_ – *F*_*L*100_.

### RNA sequencing

*Synechocystis* fast growing mutant strains as well as the control strain (sgRNA-NT0) were cultivated in photobioreactor in turbidostat mode under the same conditions as they were enriched (see above). At 3 days post induction, 40 mL culture (OD_730 nm_ = 0.2) was sampled. Cells were collected by centrifugation for 5 min at 4 °C, and total RNA was extracted immediately afterwards. Ribosomal rRNA was depleted using Illumina Ribo-Zero rRNA Removal Kit (Bacteria). Library preparation was carried out using NEBNext Ultra II Directional RNA Library Prep Kit (New England Biolabs) following manufacturers guidelines. Libraries were sequenced on Illumina NextSeq 500 System using NextSeq 500/550 High Output v2 kit (75 cycles). RNA sequencing reads were filtered and mapped to the genome as described previously^77^.

### NGS Data Analysis

#### Statistical analysis

All analyses were performed using the R programming language and are documented in R markdown notebooks available at https://m-jahn.github.io/. Data for all competition experiments performed with the sgRNA library can be accessed at https://m-jahn.shinyapps.io/ShinyLib/.

First, data tables from different sequencing runs were merged into a single master table. For simplicity, harvesting time points 12 and 30 days for the sodium chloride condition (NACL) were re-labelled as 16 and 32 days to correspond to time points of all other samples. This did not influence the calculation of generation time or fitness score, and was done only to display these samples along with corresponding L-lactate samples. The R package DESeq2 ^78^ was used to determine fold changes between conditions as well as significance metrics (multiple hypothesis adjusted p-value, Benjamini-Hochberg procedure). Determination of fold change and significance was based on aggregating 4 independent biological replicates for all cultivations. Gene-wise annotation was added based on <u>Uniprot</u> (IDs, protein properties, GO terms) and <u>CyanoBase</u> (functional categories). Altogether 7,119 unique sgRNAs corresponding to 3,541 unique genes (without non-coding RNAs) were included in the analysis. The coverage per sample in terms of quantified sgRNAs and median read count per gene is shown in Supplemental Fig. 11.

#### Unsupervised clustering

To cluster genes based on depletion/enrichment pattern, a dissimilarity matrix was computed using R’s *dist* function with distance measure *euclidean*. Clustering was performed using function *hclust* with method *ward.D2*. Silhouette analysis was performed to find the optimal number of clusters (*silhouetteAnalysis* from package *silhouette*) and showed equally good separation for 3 to 9 clusters. A number of 5 clusters was chosen representing sgRNAs with decreasing level of depletion (1-4) as well as unchanged sgRNAs (5).

#### Gene ontology (GO) enrichment

For GO term enrichment, the TopGO package by Alexa *et al*.^79^, was used to determine GO terms associated with sgRNAs for clusters 1 to 4 (TopGO method ‘Fisher, eliminating’). The resulting list of GO terms was filtered by dispensability scores obtained using REVIGO (http://revigo.irb.hr/, threshold ≤ 0.5). Furthermore, GO terms annotated with less than 5 or more than 200 unique sgRNA/genes or p-value > 0.03 were filtered out.

#### Enrichment of sgRNA mutants with increased L-lactate tolerance

To find genes involved in L-lactate tolerance, sgRNAs enriched specifically for presence of sodium L-lactate but not sodium chloride were selected (threshold: log_2_ FC ≥ 2, −log_10_ p-value ≥ 20). Of the 78 sgRNAs falling into this category, none was enriched under sodium chloride. In contrast to the few sgRNAs that were enriched, thousands of sgRNAs were depleted during growth in the presence of L-lactate. To find sgRNAs specifically depleted under L-lactate but not sodium chloride, a differential fitness score was calculated between the two conditions (*dF* = *F_NaCl_* - *F_Lac_*). The top 200 sgRNAs with highest *dF* were selected and 6 genes were found with both sgRNAs strongly depleted (see also Supplemental Fig. 8).

#### RNA seq data analysis

RNA sequencing data was initially processed as described before for NGS sequencing of library data. DESeq2 was used to determine fold changes between conditions as well as significance (4 independent biological replicates). Significant genes were selected based on the following two criteria: absolute log_2_ FC ≥ 1, adjusted p-value ≤ 0.05). Unsupervised clustering of significantly different genes based on expression in all mutants was performed as described for NGS data analysis.

### Estimation of mutant growth rates

The change in a mutant’s abundance within the total population, combined with the population average growth rate (known from the dilution rate of the turbidostat), allows estimation of the growth rate of each mutant. The rate of enrichment or depletion of a mutant (μ_*diff*_) was defined as the population growth rate (μ_*pop*_) subtracted by the mutant’s individual growth rate (μ_*mut*_).

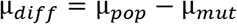

The depletion of a mutant from the library can be modeled as a function of time, where the mutant fraction *f* at time point *t* becomes:

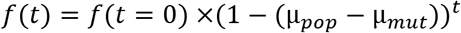

The mutant fraction *f* at different times as well as the average growth rate of the population are known parameters, allowing to estimate growth rate for all mutants.

### Correlation between gene expression variability and fitness

To correlate gene fitness (obtained from sgRNA library competition experiments) with variability in gene expression, a previously published proteomics data set was used^17^. The dataset contains mass spectrometry based protein measurements of *Synechocystis* for 2,000 proteins at 5 different growth rates, each for CO_2_ and light limitation. The variability of a protein was defined as growth-rate dependent change in abundance, and determined as p-value from analysis of variance (ANOVA). If a protein changed abundance significantly with growth rate (either up or down-regulation) a lower p-value was obtained. Proteins were binned into groups according to p-value ranges (1-0.1, 0.1-0.05, 0.05-0.01, 0.01-0.001, <0.001) and fitness score of sgRNAs associated with the respective proteins in a group were plotted (Supplemental Fig. 1).

### Absorption spectrum of L-lactate-tolerant mutants

Selected mutants as well as the control strain (sgRNA-NT0) were cultivated in 30 mL BG-11 supplemented with 100 mM sodium L-lactate in shaking flasks at 100 μmol photons m^-2^ s^-1^ continuous light, and 500 ng/mL aTc was added at the beginning of the cultivation. 0.5 mL cell culture was sampled after induction for 3 days and cells were collected by centrifugation at 5,000 xg for 10 min. Cells were then washed twice with 1 mL PBS by centrifugation and resuspension, and then resuspended in 400 μL PBS. Two times 100 μL resuspended cells were transferred to a transparent 96-well plate. Absorption spectra of cell samples were obtained in the range of 350 to 800 nm using a photospectrometer (SpectraMax M5, Molecular Devices). Absorption spectra were normalized to the reference A_720 nm_ and relative chlorophyll a, phycocyanin and carotenoid content of cells were determined using the ratio of A_680 nm_/A_720 nm_, A_630 nm_/A_720 nm_, and A_490 nm_/A_720 nm_, respectively.

### Droplet microfluidics screening of L-lactate-producing sgRNA library

The L-lactate-producing *Synechocystis* sgRNA library were grown in shake-flask in a climatic chamber. Cultures were supplemented with antibiotics (12.5 μg/mL chloramphenicol, 25 μg/mL spectinomycin) and aTc (1 μg/mL). For the productivity assay, cells were harvested (OD_720 nm_=0.4-0.6), washed to OD_720 nm_=0.15, encapsulated in droplets as previously described^69^. The droplet emulsion was incubated in an illuminated syringe (approximately 150 μE/s/m^2^) for 4 h for L-lactate secretion. The emulsion was then injected onto a picoinjection chip^80^ with 3 pL of a fluorescence-based L-lactate assay mixture (Cayman Chemical). The emulsion was injected at a flow rate of 70 μL/h,the spacer oil separated the droplets at a flow rate of 500 μL/hr and the lactate assay mixture was injected at 30 μL/h. The pico-injected emulsion was collected in a 1 mL plastic syringe protected from light. After 40 minutes of pico-injection, the collected emulsion was gently mixed and re-injected into a droplet sorting device as previously described^81^. The flow rates used were 1000 μL/h for the spacer oil and side oil, and the emulsion was injected at 100 μL/h, corresponding to a droplet sorting rate of 1.5 kHz. Sorted droplets were collected using a withdrawal rate of 1000 μL/h. Approximately 1,800,000 droplets were screened and the top 2% fluorescent were sorted, corresponding to approximately 36,000 droplets. After sorting, the collected droplet emulsion was broken with 10% v/v 1H, 1H, 2H, 2H-perfluoro-1-octanol (Sigma Aldrich), followed by centrifugation and evaporation to remove the residual liquid. Cells were resuspended in DMSO and heated at 95°C for 2 min to release genomic content. The cell lysate was used as template to directly amplify the sgRNA region using primer pair LUYA271/LUYA300. Then adaptors were incorporated into the PCR product using primer pair LUYA593/LUYA594, followed by NGS library preparation as described above.

### Droplet microfluidics NGS data filtering

Using NGS obtained on the sorted droplets, that approximately 8,000 unique sgRNAs were detected from approximately 5 million mapped reads. sgRNAs with fewer than 32 mapped reads in a sample were removed from that sample. To calculate ‘enrichment factors’ for each sgRNA, the relative abundance of the sgRNA in the sorted droplet was divided by its relative abundance in the total library before droplet encapsulation. Next, sgRNAs with a 10-fold difference in enrichment factor between replicates were removed. The remaining sgRNAs were then ranked by enrichment factor for each sorted sample. sgRNAs in the top 10% were given a score of 1 for that sample. Clones with a score 2/4, summed over all samples were determined to have potential to increase L-lactate productivity.

### L-lactate production in batch and photonfluxostat

The L-lactate-producing *Synechocystis* sgRNA library mutants were first cultivated in shakeflasks in a climatic chamber. Cultures were pre-cultivated in BG-11 supplemented with antibiotics (12.5 μg/mL chloramphenicol, 25 μg/mL kanamycin) and induced with aTc (1 μg/mL) 2 days prior to the start of the experiment. Then mutants were transferred to shake-flasks or in an 8-tube photobioreactor (see library cultivation methods) with a starting OD_720 nm_ of 0.1, supplemented with aTc and antibiotics as stated above. Cultures in the photobioreactor were grown in a ‘photonfluxostat’ mode, where light is increased based on cellular density to extend log phase. The amount of light given to the culture equals to the cellular density multiplied by a light regime factor (1000 μmol photons m^-2^ s^-1^ OD_720 nm_^-1^). L-lactate was measured using a L-lactate fluorescent kit (Cayman Chemical) according to the manufacturer’s instructions.

## Supplemental Materials

**Supplemental Data 1:** Fasta file with gene name, gene locus, protospacer, and other regions used for cloning. (.fasta)

**Supplemental Data 2**: Fasta file containing ncRNA sequences and IDs as represented in Supplemental Data 1. (.fasta)

**Supplemental Data 3**: Summary of all growth conditions, including dilution rates and generation times. (.csv)

**Supplemental Data 4:** Table of the abundance of all detected sgRNAs and their fold-change over cultivation time for all conditions. (.csv)

**Supplemental Data 5**: Table of sgRNA fitness scores in L100, L300, and LD conditions

**Supplemental Data 6**: Calculated specific growth rates for selected genes in central carbon metabolism and faster-growing mutants (.csv)

**Supplemental Data 7**: Gene expression data for *slr1916*, *ssl2892*, *sll1969*, and *slr1340* sgRNA strains compared to NT0 control strain. (.csv)

**Supplemental Data 8:** List of all clones that pass the enrichment threshold in the droplet-based lactate assays and their frequency. (.xlsx)

**Supplemental Data 9:** Comparison of *Synechocystis* gene fitness scores with gene essentiality as determined by flux balance analysis of a genome-scale model. (.xlsx)

**Supplemental Data 10**: Comparison of *Synechocystis* gene fitness scores to essentiality of *Synechoccocus* orthologs determined by Tn-Seq. (.xlsx)

## Supplemental information

Includes Supplemental Figures and Tables (.pdf)

**Supplemental Figure 1:** Highly regulated proteins have lower average fitness.

**Supplemental Figure 2:** Estimated growth rate of selected mutants.

**Supplemental Figure 3:** Comparison of fitness scores for all genes in L100 condition compared to LD condition.

**Supplemental Figure 4:** Comparison of fitness scores for all genes in L300 condition compared to LD condition.

**Supplemental Figure 5:** Fitness scores for all genes in the phylloquinone pathway.

**Supplemental Figure 6:** Time courses and functional category assignments of genes with only 1 sgRNA enriched when L-lactate was present in culture medium.

**Supplemental Figure 7:** Time-course of growth rate of re-constructed L-lactate-tolerance mutants in batch cultures.

**Supplemental Figure 8:** Selected clones that were depleted from the turbidostat during L-lactate-stress cultivation but not NaCl stress cultivation.

**Supplemental Figure 9:** Schematic of microfluidics-based sorting of L-lactate producers library. Examples of L-lactate fluorescence histograms from droplets at different time points after dCas9 induction.

**Supplemental Figure 10:** Correlation of log_2_ fold change for each sgRNA pair per gene, for all conditions.

**Supplemental Figure 11:** Median read count and coverage of samples in terms of quantified unique sgRNAs for each condition and time point.

**Supplemental Table 1**: Primers used in this study.

## Acknowledgements

We would like to thank Filipe Brancos dos Santos and Klaas Hellingwerf for helpful discussions. This work was funded by Swedish Research Council [2016-06160] and the Swedish Foundation for Strategic Research SSF [RB14-0013].

## Author contributions

K.S. and L.Y. designed and cloned the CRISPRi library, performed competition experiments and next-generation sequencing. L.Y. cloned and cultivated faster-growing mutants. K.S. cloned and cultivated lactate over-producing mutants. K.S. and S.B. performed droplet sorting. J.A.S. performed RNA-Seq data processing. M.J. performed data processing and analysis for competition experiments and RNA-Seq. All authors designed the experiments. L.Y., K.S., P.H. and M.J. wrote the manuscript. All authors read and approved the manuscript.

