## Supplemental figures and tables for "Pooled CRISPRi screening of the cyanobacterium *Synechocystis* sp. PCC 6803 for enhanced growth, tolerance, and chemical production"

for the manuscript:

### Contents

**Supplemental Figure 1.**
Highly regulated proteins have lower average fitness.

**Supplemental Figure 2.**
Estimated growth rate of selected mutants.

**Supplemental Figure 3.**
Comparison of fitness scores for all genes in L100 condition compared to LD condition.

**Supplemental Figure 4.**
Comparison of fitness scores for all genes in L300 condition compared to LD condition.

**Supplemental Figure 5.**
Fitness scores for all genes in the phylloquinone pathway.

**Supplemental Figure 6.**
Time courses and functional category assignments of genes with only 1 sgRNA enriched when L-lactate was present in culture medium.

**Supplemental Figure 7.**
Time-course of growth rate of re-constructed L-lactate-tolerance mutants in batch cultures.

**Supplemental Figure 8.**
Selected clones that were depleted from the turbidostat during L-lactate-stress cultivation but not NaCl stress cultivation.

**Supplemental Figure 9.**
Schematic of microfluidics-based sorting of lactate producers library. Examples of lactate fluorescence histograms from droplets at different time points after dCas9 induction.

**Supplemental Figure 10.**
Correlation of $\log_2$ fold change for each sgRNA pair per gene, for all conditions.

**Supplemental Figure 11.**
Median read count and coverage of samples in terms of quantified unique sgRNAs for each condition and time point.

**Supplemental Table 1**.

Primers used in this study.

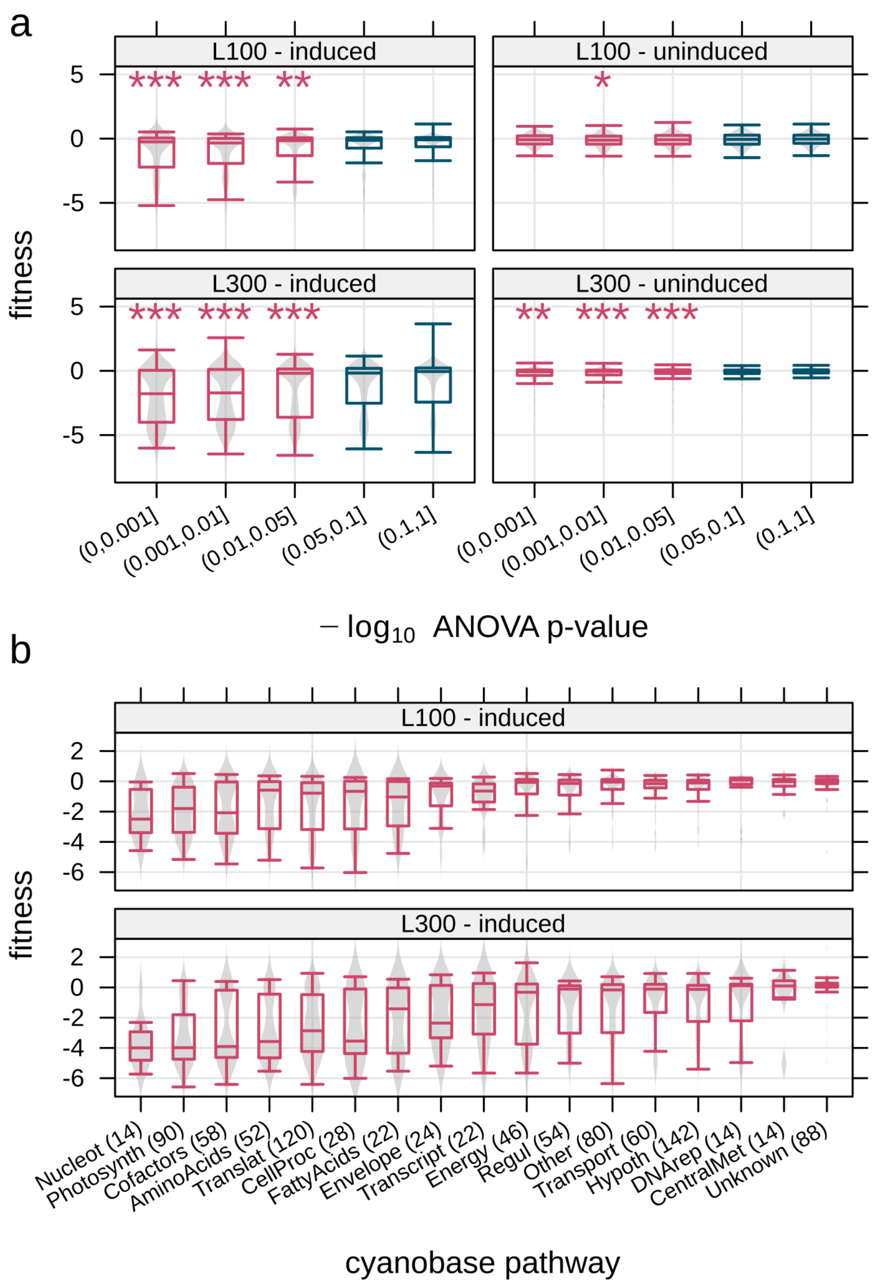


**Supplemental Figure 1. Highly regulated proteins have lower average fitness. a** Fitness score of sgRNAs broken down by protein variability. Protein variability was estimated using a proteomics study with measurement of protein abundance over several light conditions [1]. Here, the adjusted p-value from ANOVA over all light conditions was used as a metric for variability. For this comparison every sgRNA was mapped to its corresponding protein. Red, proteins with ANOVA *p-value* ≤ 0.05. Blue, proteins with ANOVA *p-value* > 0.05. Symbols, significance of distribution being different from last group (0.1-1) according to Student's t-test: '*' *p-value* ≤ 0.05, '**' *p-value* ≤ 0.01, '***' *p-value* ≤ 0.001 **b** Fitness score of sgRNAs associated with most significantly changing proteins (*p-value* ≤ 0.05, red in **a**), broken down by cyanobase pathways. In brackets, number of unique sgRNAs per pathway.

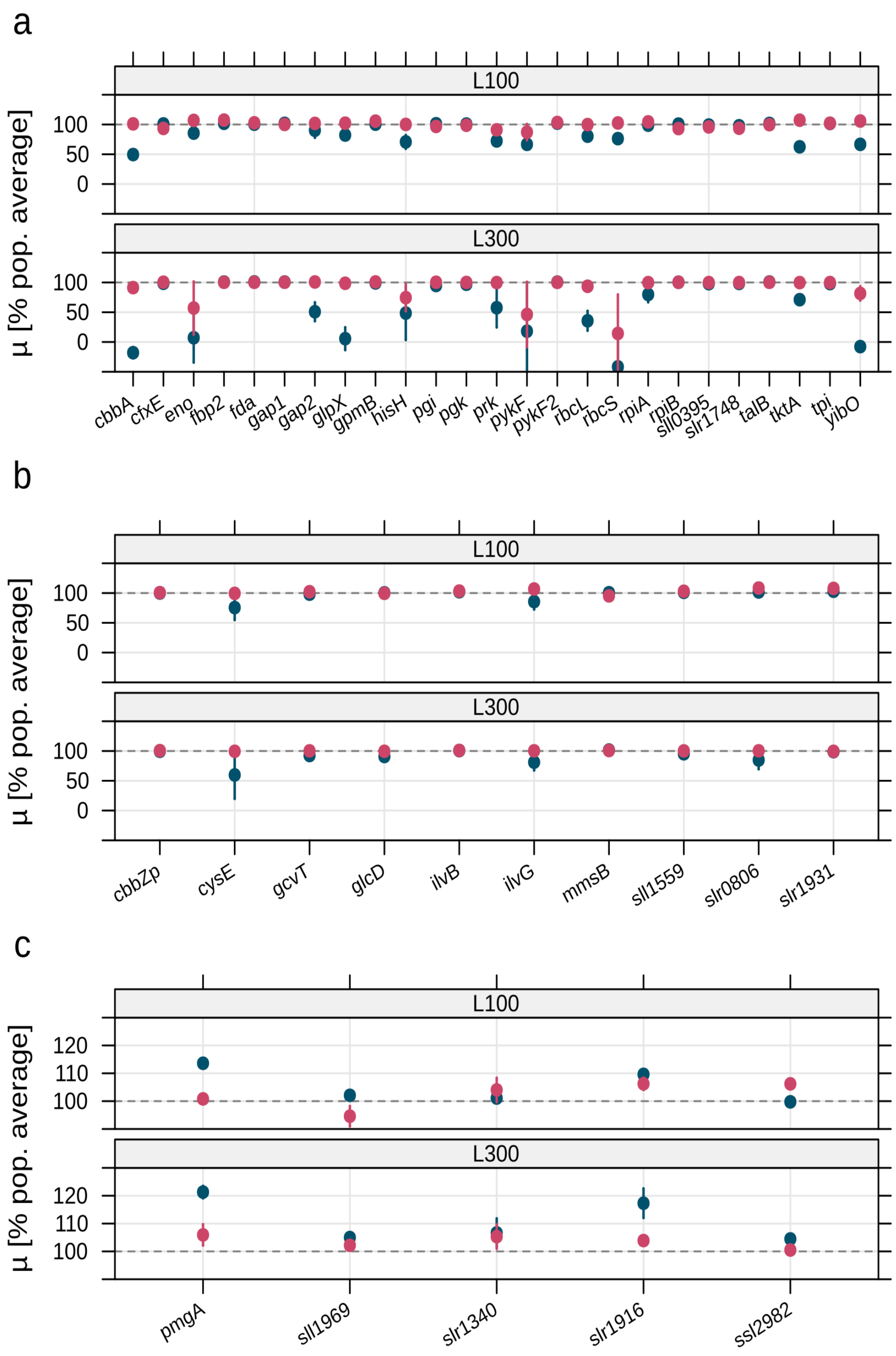


**Supplemental Figure 2. Estimated growth rate of selected mutants.** Mutant growth rate was determined from depletion of sgRNAs over time as described in Methods (mean and standard deviation of two sgRNAs per gene). Growth rate is displayed as % of population growth rate. Red - uninduced control, blue - induced culture. **a** Selected genes from the Calvin cycle. **b** Selected genes for photorespiration. **c** Selected faster growing mutants.

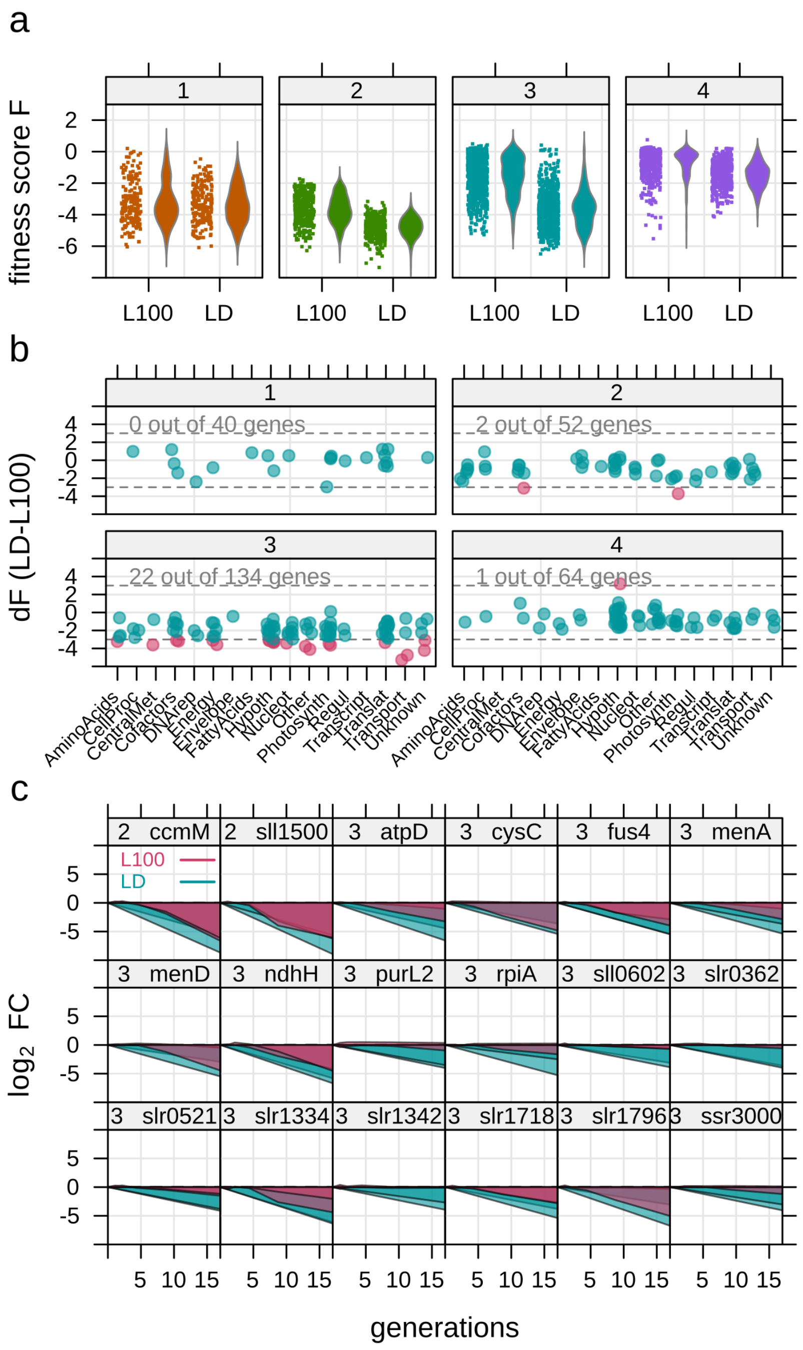


**Supplemental Figure 3. Genes with condition-dependent fitness for L100 and LD. a** Distribution of fitness score *F* for all sgRNAs in cluster 1 to 4. Fitness score indicates the degree of enrichment (positive) or depletion (negative) of an sgRNA. Comparison between low light (L100 - light with 100 µmol photons $m^{-2}$ $s^{-1}$) and light-dark cycle (LD - light with 0-300 µmol photons $m^{-2}$ $s^{-1}$). **b** Difference between fitness score of L100 and LD (d*F*) for genes with both sgRNAs in the same cluster. Differentially depleted/enriched sgRNAs indicated in red, threshold: 3 <= dF <= -3. **c** Selection of top 18 genes with d*F* above threshold as described in **b**. Only genes with 2 depleted/enriched sgRNAs and annotated description were selected.

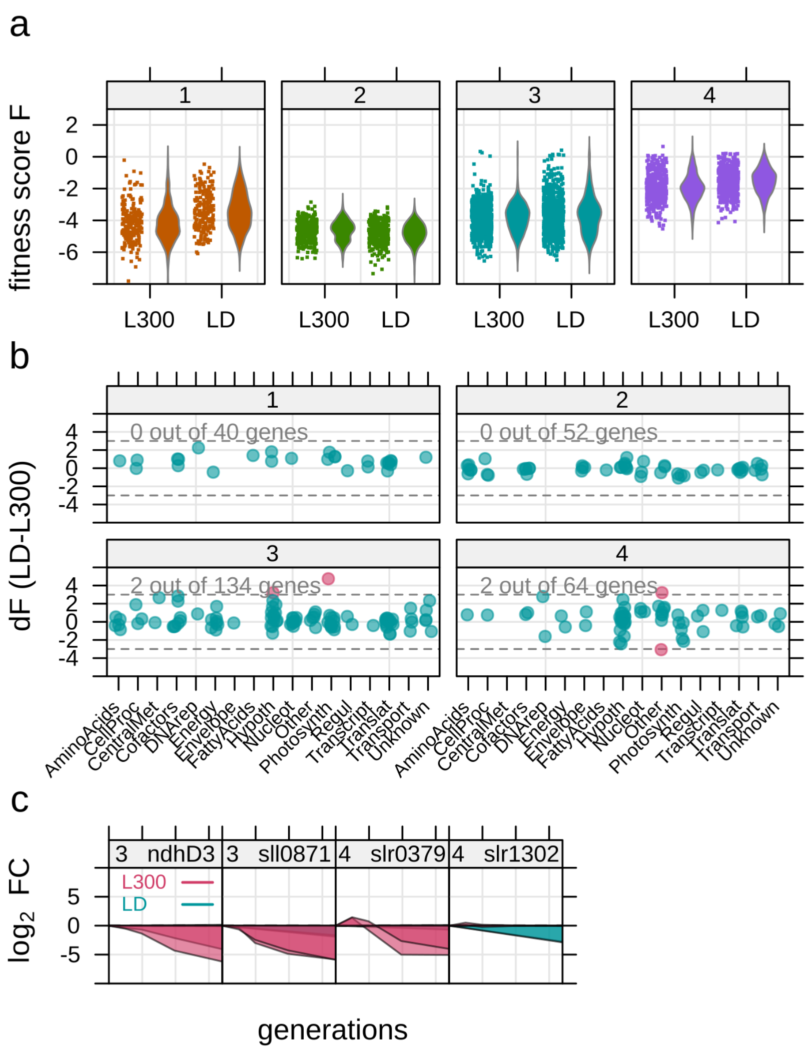


**Supplemental Figure 4. Genes with condition-dependent fitness for L300 and LD. a** Distribution of fitness score *F* for all sgRNAs in cluster 1 to 4. Fitness score indicates the degree of enrichment (positive) or depletion (negative) of an sgRNA. Comparison between high light (L300 - light with 300 µmol photons $m^{-2}$ $s^{-1}$) and light dark cycle (LD - light with 0-300 µmol photons $m^{-2}$ $s^{-1}$). **b** Difference between fitness score of L300 and LD (d*F*) for genes with both sgRNAs in the same cluster. Differentially depleted/enriched sgRNAs indicated in red, threshold: 3 <= dF <= -3. **c** All genes with d*F* above threshold as described in **b**. Only genes with 2 depleted/enriched sgRNAs and annotated description were selected.

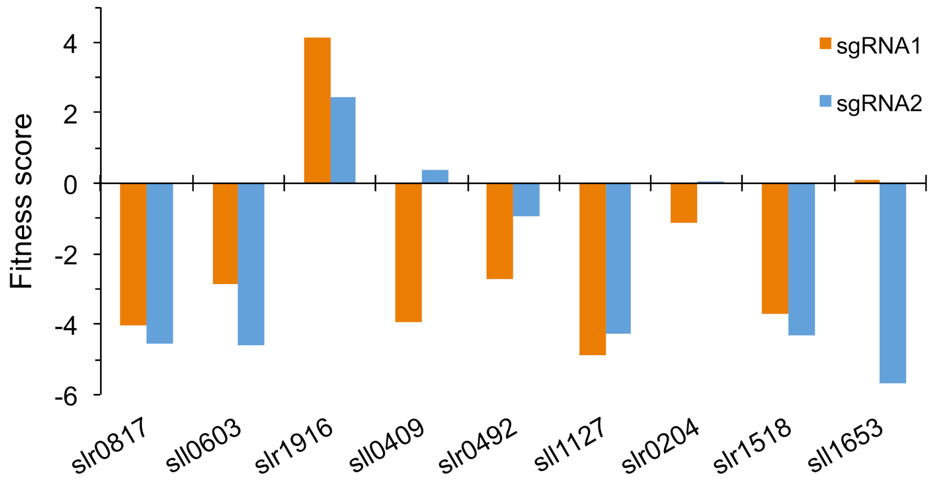


**Supplemental Figure 5. Fitness score of sgRNAs targeting phylloquinone biosynthetic pathway genes.** *slr0817*: isochorismate synthase (MenF); *sll0603*: SEPHCHC synthase (MenD); *slr1916*: (*1R,6R*)-2- succinyl-6-hydroxy-2,4-cyclohexadiene-1-carboxylic acid synthase (MenH); *sll0409*: *o*-succinylbenzoate synthase (MenC); *slr0492*: *o*- succinylbenzoate-CoA ligase (MenE); *sll1127*: 1,4-dihydroxy-2-naphthoate synthase (MenB); *slr0204*: 1,4-dihydroxy-2-naphthoate-CoA thioesterease; *slr1518*: 1,4-dihydroxy-2-naphthoate prenyltransferase (MenA); *sll1653*: demethylmenaquinone/demethylphylloquinone methyltransferase (MenG).

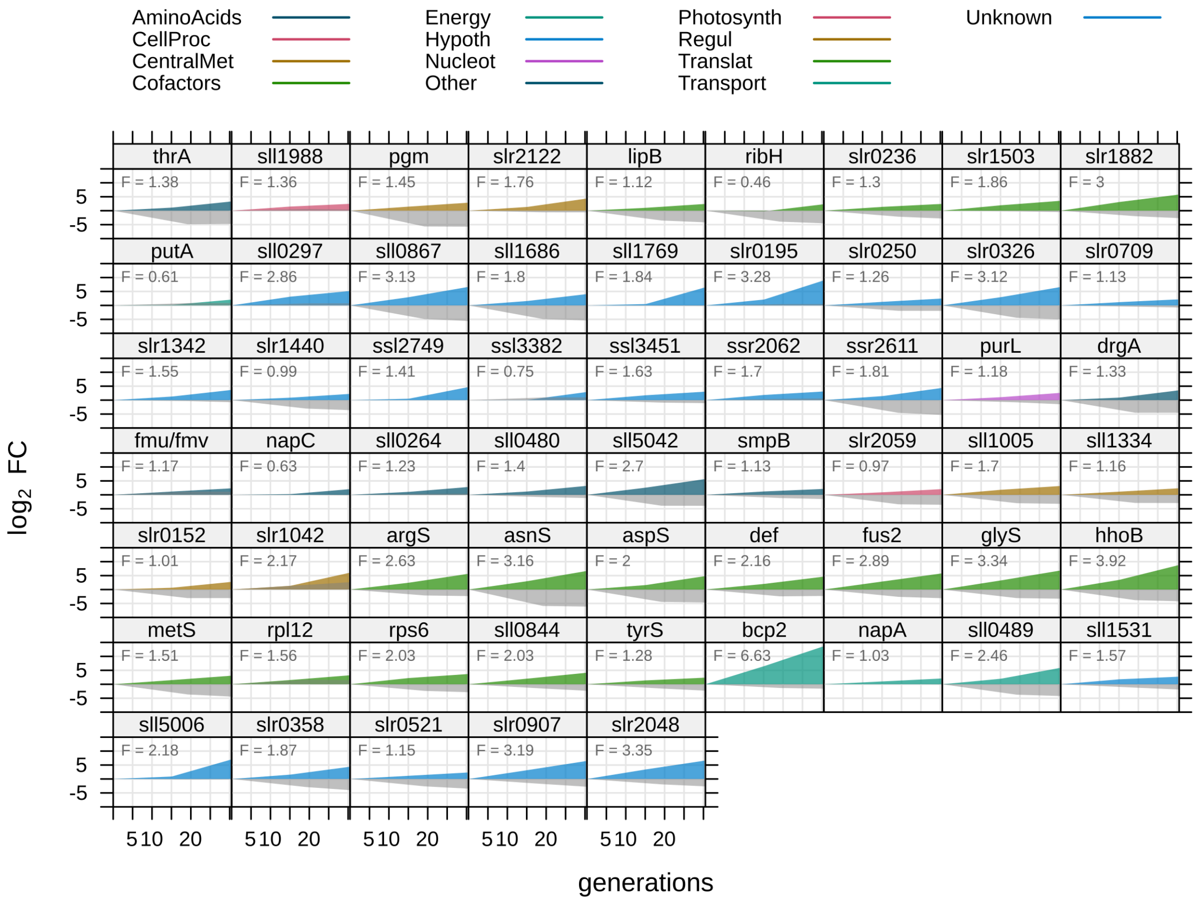


**Supplemental Figure 6. Genes with only 1 sgRNA enriched when L-lactate was present in culture medium.** Depicted is the $\log_2$ fold change ($\log_2$ FC) over number of generations the cells were cultivated in a turbidostat bioreactor. Colored area: sgRNA in presence of 0.1 M L-lactate, color-coded by cyanobase functional category, grey area: sgRNA in presence of 0.1 M sodium chloride. *F*, fitness score for sgRNA in presence of lactate. Fitness score is positive for enrichment and negative for depletion of sgRNAs.

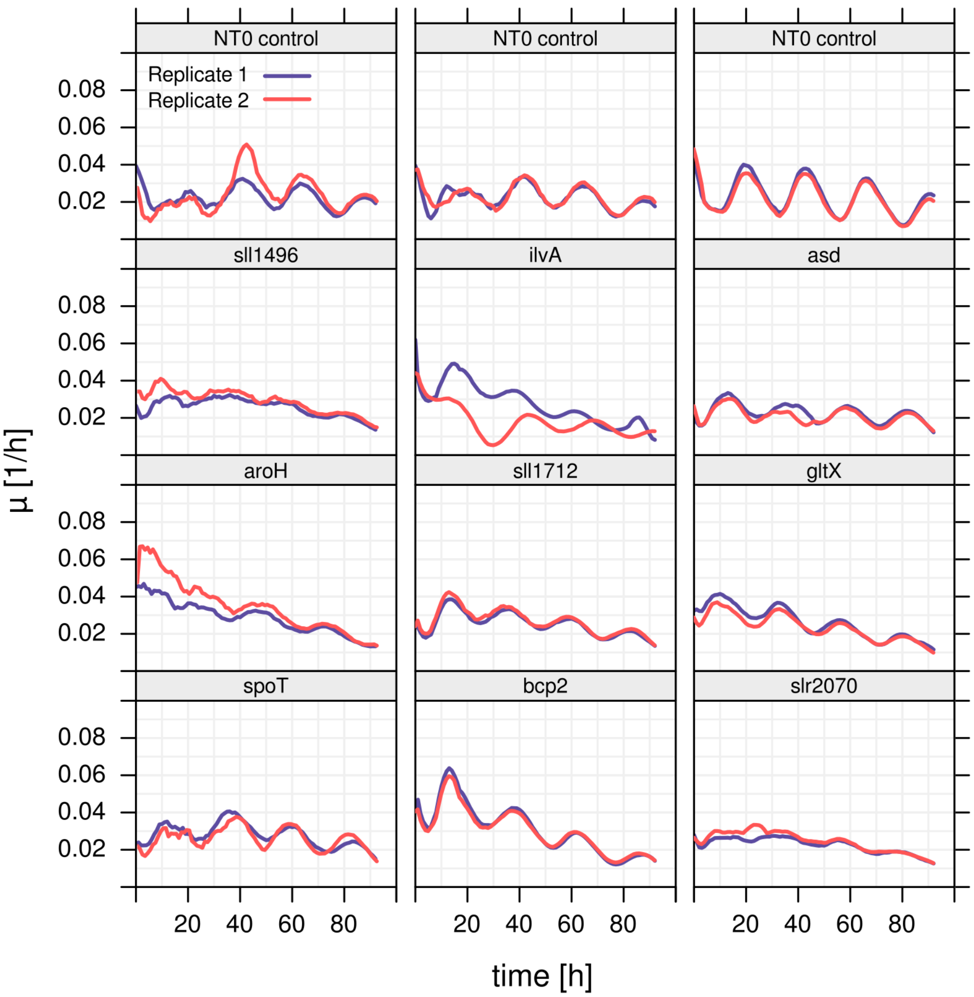


**Supplemental Figure 7. Cultivation of L-lactate tolerant sgRNA mutants for initial 100 hours.** Selected sgRNA mutants were reconstructed and grown in axenic turbidostat cultivations in presence of 0.1 M L-lactate (n = 2). Cultivation conditions for all strains were 100 µmol photons $m^{-2}$ $s^{-1}$ and 1% $CO_2$. The control strain (NT0) contains an sgRNA with no target site in *Synechocystis* genome. Specific growth rate µ over time was calculated in a sliding window of step width 5 hours. The oscillation in growth rate is caused by the inherent circadian rhythm.

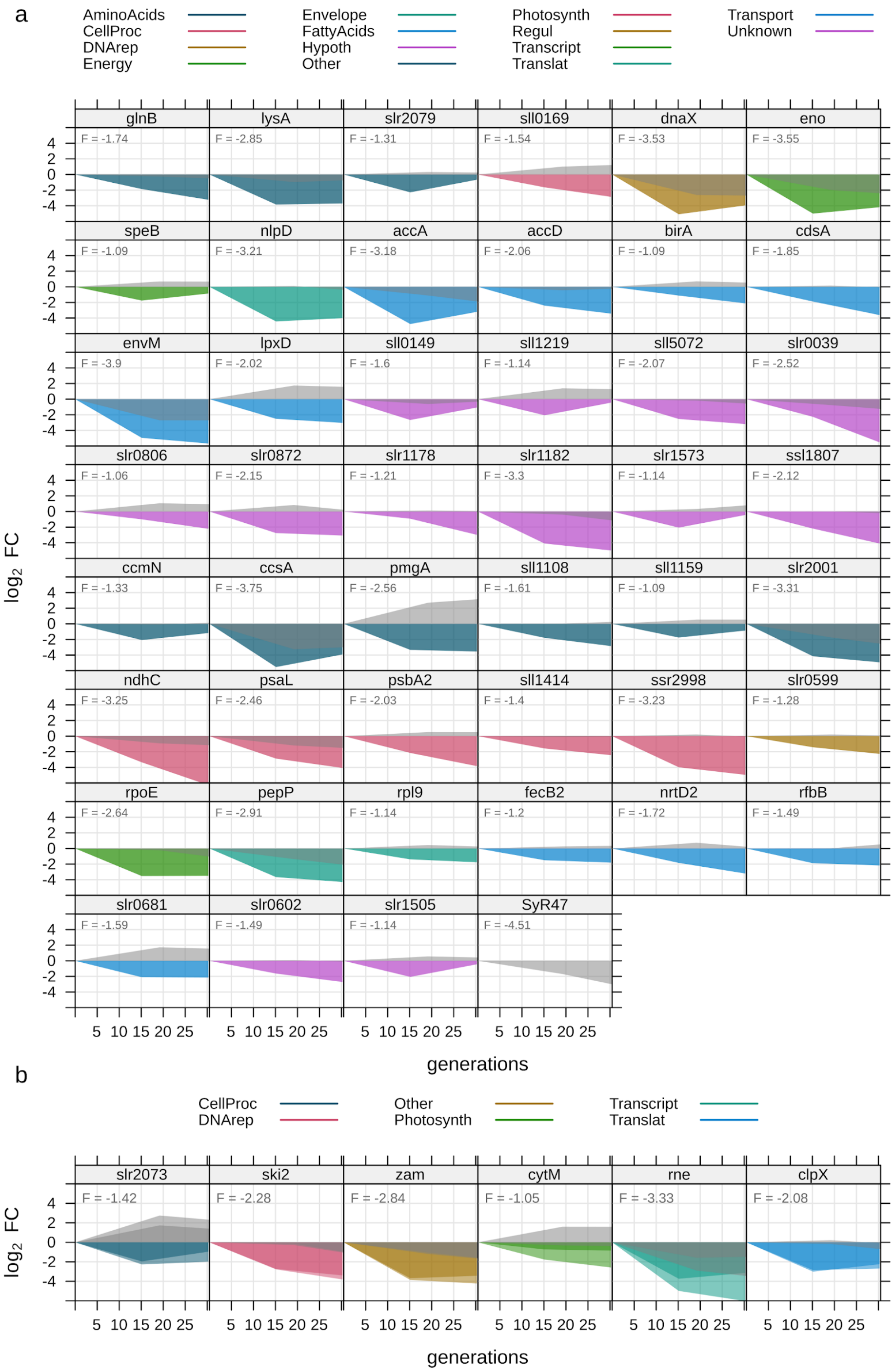


**Supplemental Figure 8. Genes that were depleted when L-lactate was present in culture medium.** **a** Genes where only 1 sgRNA was depleted. **b** Genes where both sgRNAs were depleted. Depicted is the $\log_2$ fold change ($\log_2$ FC) over number of generations when the cells were cultivated in a turbidostat bioreactor. Colored area: sgRNA in presence of 0.1 M L-lactate, color-coded by cyanobase functional category, grey area: sgRNA in presence of 0.1 M sodium chloride. *F*, fitness score for sgRNA in presence of L-lactate. Fitness score is positive for enrichment and negative for depletion of sgRNAs.

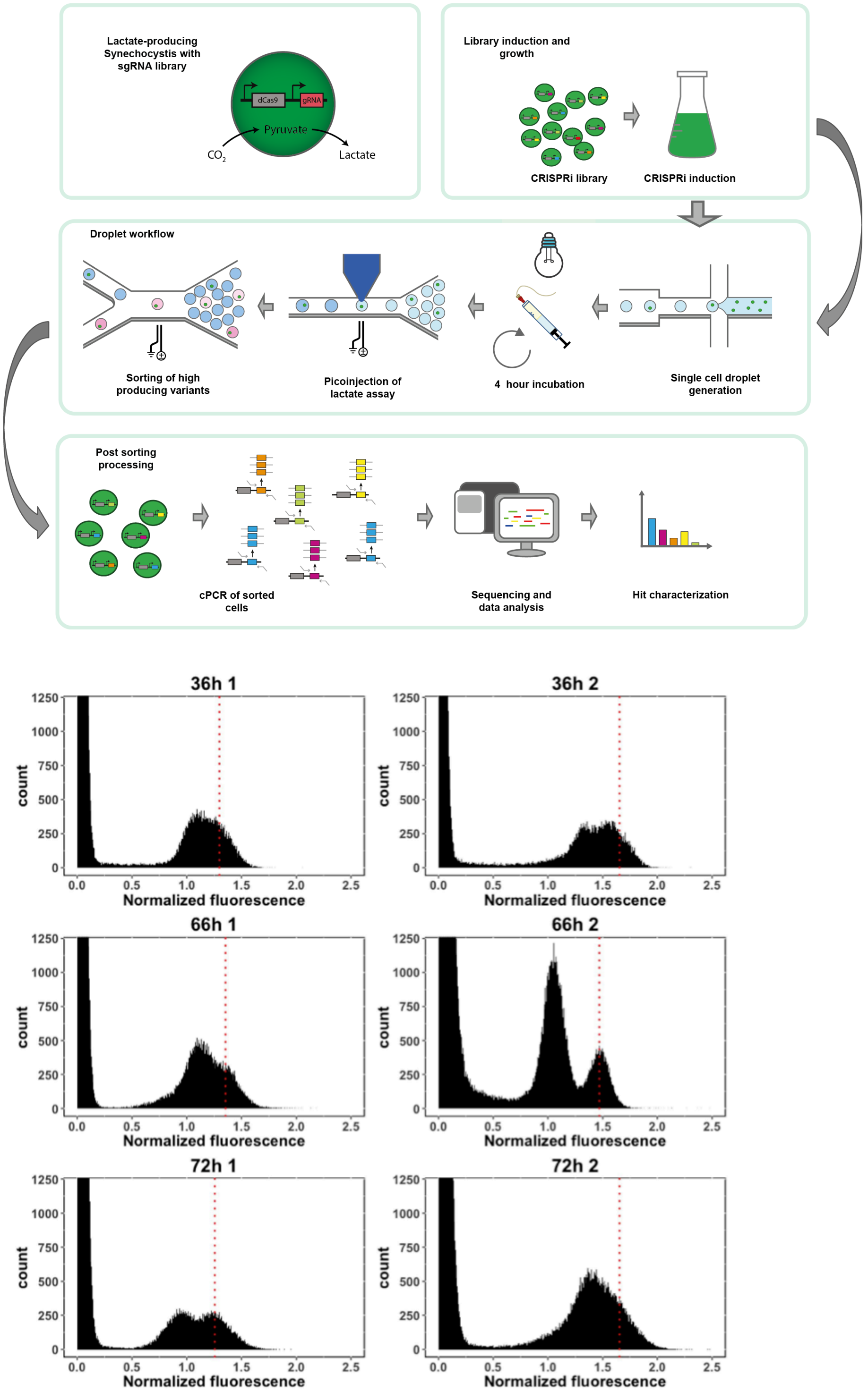


**Supplemental Figure 9. Method details on droplet sorting. a** Schematic of microfluidics-based sorting of lactate producers library. **b** Histogram of lactate fluorescence intensity from droplets at different time points after dCas9 induction. Dotted line - threshold between low (left) and high (right, sorted droplets) fluorescent sub-populations.

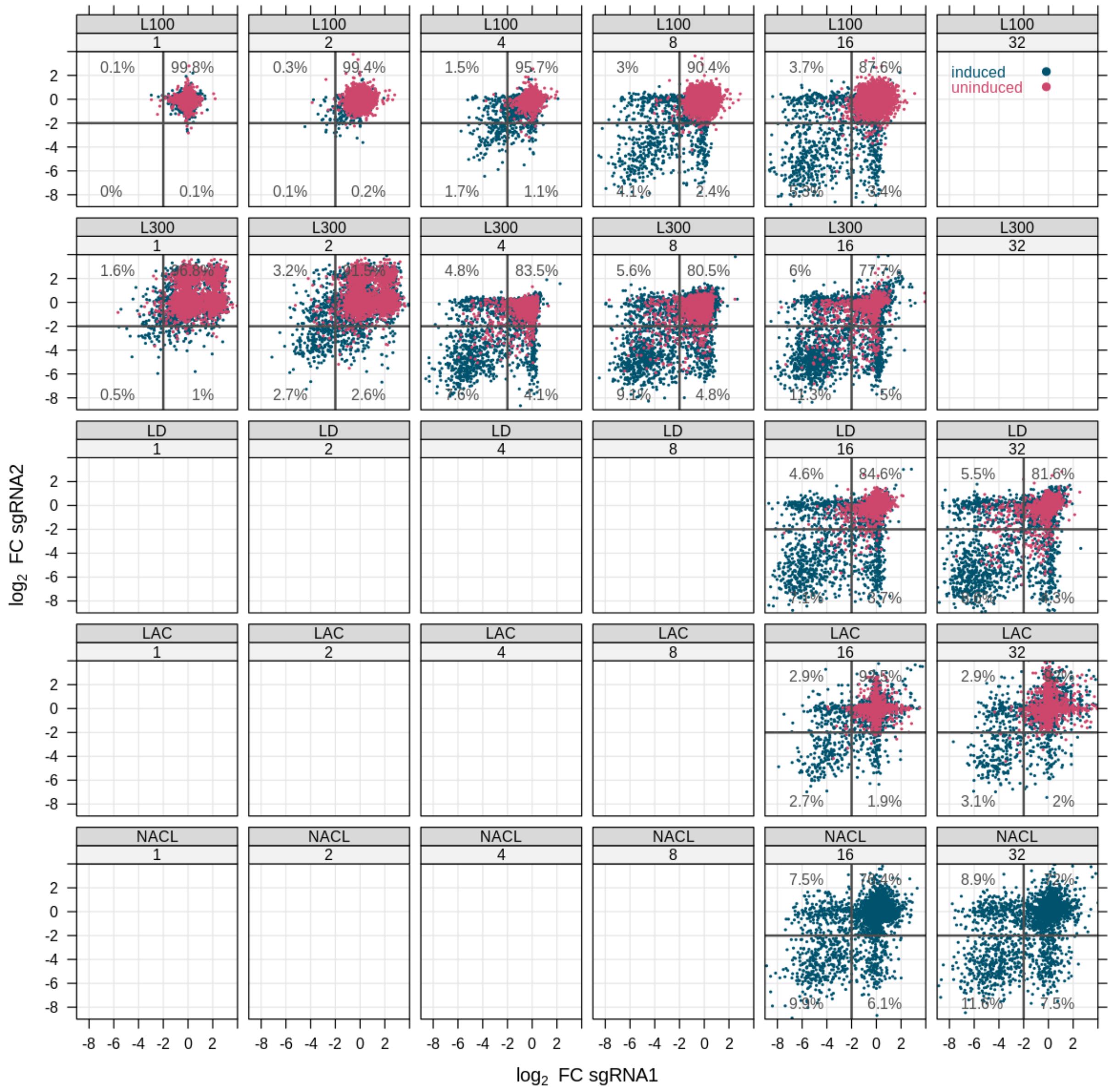


**Supplemental Figure 10. Correlation of sgRNA pairs for each gene.** Correlation of the $\log_2$ FC of sgRNA1 *versus* sgRNA 2 for each gene, broken down by condition. Each panel shows an overlay of induced (blue) and uninduced (red) samples. Panel numbers indicate time in days. L100 - light with 100 µmol photons $m^{-2}$ $s^{-1}$, L300 - light with 100 µmol photons $m^{-2}$ $s^{-1}$, LD - light-dark cycle, LAC - addition of 0.1 M L-lactate, NACL - addition of 0.1 M sodium chloride.

**a** **b**

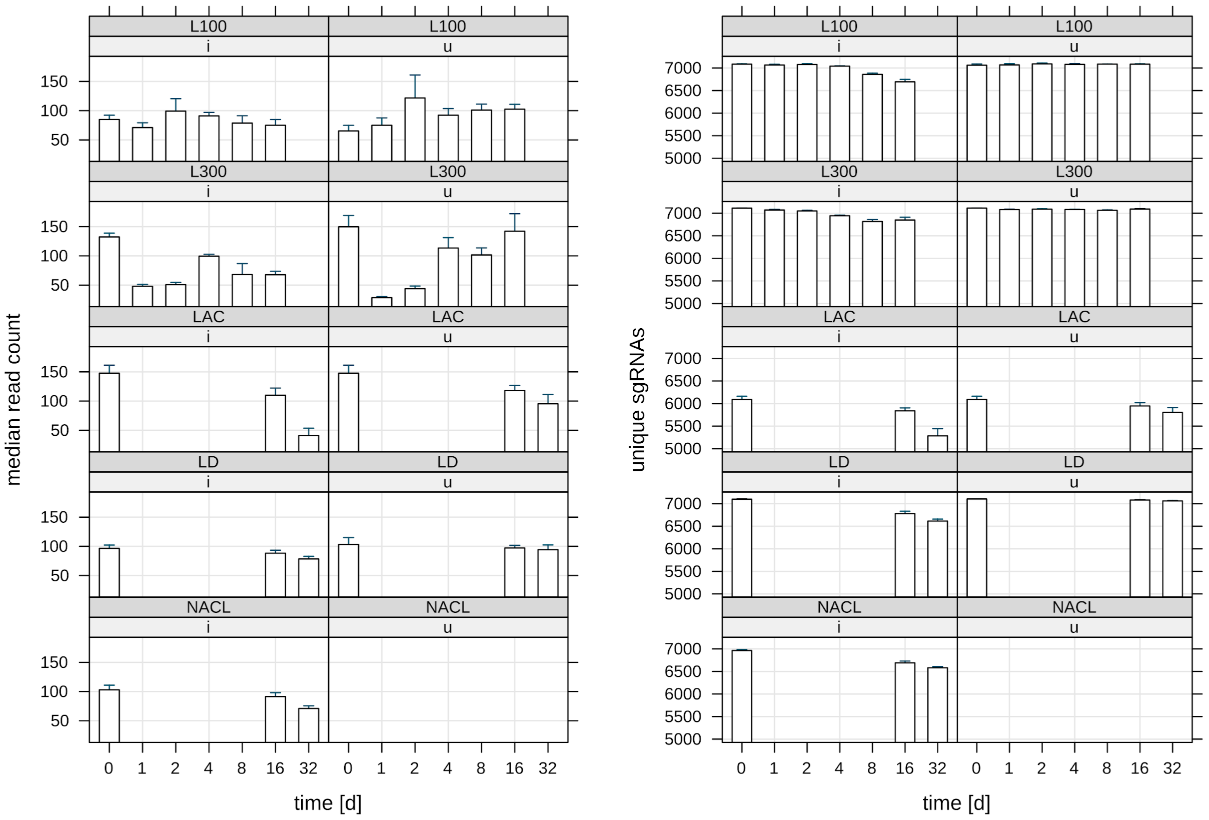


**Supplemental Figure 11. a** Median read count per condition. Bars and whiskers correspond to mean and standard deviation of 4 biological replicates. **b** Coverage of samples in terms of quantified unique sgRNAs per condition and time point. Bars and whiskers correspond to mean and standard deviation of 4 biological replicates. L100 - light with 100 µmol photons m$^{-2}$ s$^{-1}$, L300 - light with 300 µmol photons m$^{-2}$ s$^{-1}$, LD - light-dark cycle, LAC - addition of L-lactate, NACL - addition of sodium chloride.

**Supplemental Table 1. Primers used in this study**

| Primer | Sequence | Usage |
|---|---|---|
| LUYA271 | ATCTCAGGGGGAATCATTGCTTT | verify integration of sgRNA into neutral site |
| LUYA300 | TGAGCAGAATTGGGAAGGACGT | verify integration of sgRNA into neutral site |
| LUYA503 | GCTGCAATGATACCGCGAGAACCACGCTCACCGGCTCCA | mutate BsaI site on neutral site vector |
| LUYA504 | TGGAGCCGGTGAGCGTGGTTCTCGCGGTATCATTGCAGC | mutate BsaI site on neutral site vector |
| LUYA505 | AGAGGTCTCCAGTGATAGAGATACTGGGA | add BsaI overhangs to sgRNA library oligos |
| LUYA506 | AGAGGTCTCATTTTAACTTGCTATTTCTAGCT | add BsaI overhangs to sgRNA library oligos |
| LUYA507 | CTAGGTCTCTCACTGATAGGGATGTCAATCT | add BsaI overhangs to neutral site vector |
| LUYA508 | AGTGGTCTCTAAAATAAGGCTAGTCCGTTATCA | add BsaI overhangs to neutral site vector |
| LUYA530 | CCTACGTCTCTCAGTGATAGAGATACTGGGAGCTA | add Esp3I overhangs to sgRNA library oligos |
| LUYA531 | AGATCGTCTCGCCTTATTTTAACTTGCTATTTCTAGCTCTA AAAC | add Esp3I overhangs to sgRNA library oligos |
| LUYA532 | CGTATTTCGACTCGCTCAGGCGCAATCAC | mutate Esp3I site on neutral site vector |
| LUYA533 | GCCTGAGCGAGTCGAAATACGCGATCGCTGTTAAAAG | mutate Esp3I site on neutral site vector |
| LUYA534 | CTATCGTCTCCACTGATAGGGATGTCAATCTC | add Esp3I overhangs to neutral site vector |

| | | |
|---|---|---|
| LUYA535 | AAAACGTCTCTAAGGCTAGTCCGTTATCAACTT | add Esp3I overhangs to neutral site vector |
| LUYA593 | ACACTCTTTCCCTACACGACGCTCTTCCGATCTCAGTGATAGAGATACTGGGAGC | 1st step PCR for NGS |
| LUYA594 | GACTGGAGTTCAGACGTGTGCTCTTCCGATCTGCCTTATTTTAACTTGCTATTTCTAG | 1st step PCR for NGS |
| LUYA605 | CTATCGCCTGGGAGGCCTGAATGTGCGTTTTAGAGCTAGAAATAGCAAGTTA | *sll1968* sgRNA construction |
| LUYA606 | TAAAACGCACATTCAGGCCTCCCAGGCGATAGCTCCCAGTATCTCTATCACT | *sll1968* sgRNA construction |
| LUYA647 | CTATCAGAGCCAGATTGTTGATAGGTTTTAGAGCTAGAAATAGCAAGTTA | *slr1916* sgRNA construction |
| LUYA648 | TAAAACCTATCAACAATCTGGCTCTGATAGCTCCCAGTATCTCTATCACT | *slr1916* sgRNA construction |
| LUYA649 | CTAATTCTTCTCCTCCTCCCATGTTTTAGAGCTAGAAATAGCAAGTTA | *slr1340* sgRNA construction |
| LUYA650 | TAAAACATGGGAGGAGGAGAAGAATTAGCTCCCAGTATCTCTATCACT | *slr1340* sgRNA construction |
| LUYA651 | CTACTTGATGGCCAATATGGGCCGTTTTAGAGCTAGAAATAGCAAGTTA | *slr1299* sgRNA construction |
| LUYA652 | TAAAACGGCCCATATTGGCCATCAAGTAGCTCCCAGTATCTCTATCACT | *slr1299* sgRNA construction |
| LUYA653 | CTAACACAACAGGATGACGGTCGTTTTAGAGCTAGAAATAGCAAGTTA | *sll1969* sgRNA construction |
| LUYA654 | TAAAACGACCGTCATCCTGTTGTGTTAGCTCCCAGTATCTCTATCACT | *sll1969* sgRNA construction |
| LUYA655 | CTAGCTCTTCGGAACGATAAATAATGGTTTTAGAGCTAGAAATAGCAAGTTA | *ssl2982* sgRNA construction |

| | | |
|---|---|---|
| LUYA656 | TAAAACCATTATTTATCGTTCCGAAGAGCTAGCTCCCAGTATCTCTATCACT | *ssl2982* sgRNA construction |
| LUYA725 | TAAAACCCGGTGGTTCCACTTCCAAGCGTAGCTCCCAGTATCTCTATCACT | *asd* sgRNA construction |
| LUYA726 | CTACGCTTGGAAGTGGAACCACCGGGTTTTAGAGCTAGAAATAGCAAGTTA | *asd* sgRNA construction |
| LUYA731 | TAAAACGGTTTACGATGTGGCGATCGAAACTAGCTCCCAGTATCTCTATCACT | *ilvA* sgRNA construction |
| LUYA732 | CTAGTTTCGATCGCCACATCGTAAACCGTTTTAGAGCTAGAAATAGCAAGTTA | *ilvA* sgRNA construction |
| LUYA737 | TAAAACAAACAGTAAACTTGCCATAGGAACTAGCTCCCAGTATCTCTATCACT | *slr2070* sgRNA construction |
| LUYA738 | CTAGTTCCTATGGCAAGTTTACTGTTTGTTTTAGAGCTAGAAATAGCAAGTTA | *slr2070* sgRNA construction |
| LUYA739 | TAAAACACCCCCACCATTCACACCACCTAGCTCCCAGTATCTCTATCACT | *spoT* sgRNA construction |
| LUYA740 | CTAGGTGGTGTGAATGGTGGGGGTGTTTTAGAGCTAGAAATAGCAAGTTA | *spoT* sgRNA construction |
| LUYA729 | TAAAACCTAGCCCCACTGGTAATCTCCATAGCTCCCAGTATCTCTATCACT | *gltX* sgRNA construction |
| LUYA730 | CTATGGAGATTACCAGTGGGGCTAGGTTTTAGAGCTAGAAATAGCAAGTTA | *gltX* sgRNA construction |
| LUYA733 | TAAAACTGACTTGCGATCTGCCCAAGCCCATAGCTCCCAGTATCTCTATCACT | *sll1496* sgRNA construction |
| LUYA734 | CTATGGGCTTGGGCAGATCGCAAGTCAGTTTTAGAGCTAGAAATAGCAAGTTA | *sll1496* sgRNA construction |
| LUYA723 | TAAAACAGATGCGGTCTGTGAGCTACTAGCTCCCAGTATCTCTATCACT | *aroH* sgRNA construction |

| | | |
|---|---|---|
| LUYA724 | CTAGTAGCTCACAGACCGCATCTGTTTTAGAGCTAGAAATAGCAAGTTA | *aroH* sgRNA construction |
| LUYA727 | TAAAACGAGTTGGATCAACCTGCGCCCCTAGCTCCCAGTATCTCTATCACT | *bcp2* sgRNA construction |
| LUYA728 | CTAGGGGCGCAGGTTGATCCAACTCGTTTTAGAGCTAGAAATAGCAAGTTA | *bcp2* sgRNA construction |
| KISH064 | GCTAGGGCTTTTTCTAATCTTTTACGTTTTAGAGCTAGAAATAGCAAGTTA | *gdhA_25* sgRNA construction |
| KISH065 | AACGTAAAAGATTAGAAAAAGCCCTAGCTCCCAGTATCTCTATCACT | *gdhA_25* sgRNA construction |
| KISH080 | GCTATTTAGCGGCGGGGACTCCGGCTGTTTTAGAGCTAGAAATAGCAAGTTA | *gltA_41* sgRNA construction |
| KISH081 | GCTATAAACCCTTTAACTCATTGGTTTTAGAGCTAGAAATAGCAAGTTA | *gltA_41* sgRNA construction |
| KISH302 | GCTAGTGGAGGGCAATCGCCTTACCGGTTTTAGAGCTAGAAATAGCAAGTTA | *ssr2016_62* sgRNA construction |
| KISH303 | AACCGGTAAGGCGATTGCCCTCCACTAGCTCCCAGTATCTCTATCACT | *ssr2016_62* sgRNA construction |
| KISH304 | GCTAGGGGGTGAAGACCTTCGTGGCAGTTTTAGAGCTAGAAATAGCAAGTTA | *gltB_3* sgRNA construction |
| KISH305 | AACTGCCACGAAGGTCTTCACCCCCTAGCTCCCAGTATCTCTATCACT | *gltB_3* sgRNA construction |
| KISH306 | GCTAGGGAGACTGTTGCGGTTTTGTTTTAGAGCTAGAAATAGCAAGTTA | *sdhB_26* sgRNA construction |
| KISH307 | AACAAAACCGCAACAGTCTCCCTAGCTCCCAGTATCTCTATCACT | *sdhB_26* sgRNA construction |
| KISH308 | GCTAGGTGCCATGGCCTATACCATTGGTTTTAGAGCTAGAAATAGCAAGTTA | *atpA_asRNA_43* sgRNA construction |

| | | |
|---|---|---|
| KISH309 | AACCAATGGTATAGGCCATGGCACCTAGCTCCCAGTATCTCTATCACT | *atpA_asRNA_43* sgRNA construction |
| KISH310 | GCTAGGCGTTACGAATTTTGGCCGTTTTAGAGCTAGAAATAGCAAGTTA | *trxA2_5* sgRNA construction |
| KISH311 | AACGGCCAAAATTCGTAACGCCTAGCTCCCAGTATCTCTATCACT | *trxA2_5* sgRNA construction |
| KISH312 | GCTACCATCGCACTGGCTATGTGGTTTTAGAGCTAGAAATAGCAAGTTA | *sll1906_asRNA_43* sgRNA construction |
| KISH313 | AACCACATAGCCAGTGCGATGGTAGCTCCCAGTATCTCTATCACT | *sll1906_asRNA_43* sgRNA construction |
